# Interpreting the Evolutionary Echoes of a Protein Complex Essential for Inner-Ear Mechanosensation

**DOI:** 10.1101/2022.01.23.477425

**Authors:** Collin R. Nisler, Yoshie Narui, Deepanshu Choudhary, Jacob D. Bowman, Vincent J. Lynch, Marcos Sotomayor

## Abstract

The sensory epithelium of the inner ear, found in all extant lineages of vertebrates, has been subjected to over 500 million years of evolution, resulting in the complex inner ear of modern vertebrates. Inner-ear adaptations are as diverse as the species in which they are found, and such unique anatomical variations have been well studied. However, the evolutionary details of the molecular machinery that are required for hearing are less well known. Two molecules that are essential for hearing in vertebrates are cadherin-23 and protocadherin-15, proteins whose interaction with one another acts as the focal point of force transmission when converting sound waves into electrical signals that the brain can interpret. This interaction exists in every lineage of vertebrates, but little is known about the structure or mechanical properties of these proteins in most non-mammalian lineages. Here, we use various techniques to characterize the evolution of this protein interaction. Results show how evolutionary sequence changes in this complex affect its biophysical properties both in simulations and experiments, with variations in interaction strength and dynamics among extant vertebrate lineages. Evolutionary simulations also characterize how the biophysical properties of the complex in turn constrain its evolution and provide a possible explanation for the increase in deafness-causing mutants observed in cadherin-23 relative to protocadherin-15. Together, these results suggest a general picture of tip-link evolution in which selection acted to modify the tip-link interface, while subsequent neutral evolution combined with varying degrees of purifying selection drove additional diversification in modern tetrapods.

## INTRODUCTION

The sensory epithelium of the ear, found in all extant lineages of vertebrates, is a truly ancient adaptation. A homologous structure can be found in hagfishes (Coffin et al. 2004), the sister group to vertebrates, indicating that this specialized organ was present during vertebrate radiation and evolution that resulted in a greater capacity to integrate external stimuli. The transition to a land based lifestyle with the emergence of tetrapods was followed by a second major leap in information processing through the evolution of the middle and auditory inner ear in mammals, archosaurs (birds and crocidilians), and lepidosaurs (lizards and snakes) (fig. 1A) (Clack 2002; Fekete and Wu 2002; Coffin et al. 2004; Smotherman and Narins 2004). The ability to detect sound waves in air and with increased sensitivities would have imposed a significant evolutionary advantage on these early tetrapods, leading to a relatively rapid radiation in inner ear morphology and adaptation (Clack 2002; Coffin et al. 2004; Manley and Clack 2004). In addition to modifications made to morphological features of the middle and inner ear throughout vertebrate evolution, the molecular machinery underlying this process presumably underwent a similarly extensive series of atomic level modifications (Bankoff et al. 2017; Trigila et al. 2021). However, the aversion to fossilization exhibited by biological macromolecules coupled with a relative lack of structural and biochemical information of non-mammalian molecules involved in sound detection leaves many questions regarding the molecular evolution of this highly conserved process unanswered.

**Fig. 1.**
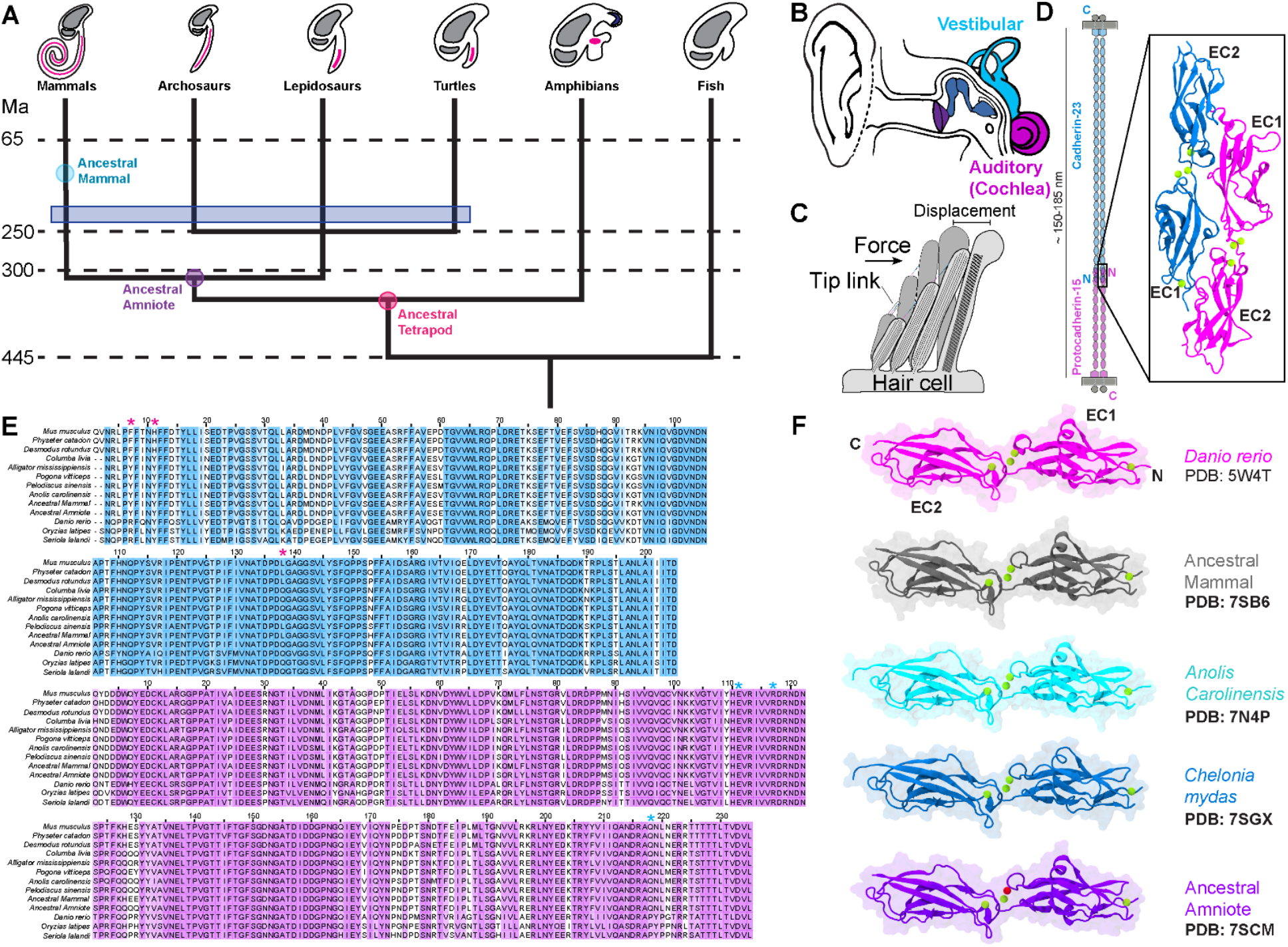
Inner ear and tip link structures and evolution. (*A*) A simple phylogeny of vertebrates, with approximate time points indicated by dashed lines, and ancestral species indicated in colored circles. The blue box represents the approximate, independent development of a tympanic middle ear in the four amniote groups. At the leaves of each branch is a cartoon representation of the inner ear for each group. Shown in grey are vestibular structures, which are an ancestral trait, and shown in pink are the auditory papillae. During vertebrate evolution, this structure elongated and diversified. (*B*) Cartoon schematic of the human ear, showing the tympanic membrane (dark purple), middle ear bones (dark blue), vestibular system (light blue), and cochlea (pink). (*C*) The stereocilia are displaced in the direction of the applied force. The tip link connects two adjacent stereocilia and pulls the cell membrane of the lower stereocilium. (*D*) The tip link is composed of a heterotetramer of CDH23 and PCDH15, which interact through their two most N-terminal EC repeats, shown in the inset. (*E*) Sequence alignments of CDH23 and PCDH15 EC1-2 for relevant vertebrate species. Pink asterisks in CDH23 indicate significant interactions that differ between vertebrate groups in the interface that were shown to be relevant in simulations, and blue asterisks in PCDH15 indicate their interaction partners. For further details on ancestral sequences, see source data 1 and 2. (*F*) Crystal structures of CDH23 EC1-2 solved for the indicated species shown in cartoon representation with overlaid transparent molecular surfaces. Bold PDB codes indicate new structures reported in this work.

All vertebrate lineages have retained the vestibular system of the inner ear derived from their common aquatic ancestor. This structure is composed of the otolith organs and semicircular canals (fig. 1A and B), which parse gravitational and acceleration information and relay these signals to the brain, allowing the animal to sense acceleration and rotation (Day and Fitzpatrick 2005). The vestibular system of fish functions in a similar manner and is believed to be homologous to that in the mammalian inner ear (Fritzsch et al. 2002; Manley et al. 2004; Fritzsch and Straka 2014). During vertebrate evolution, likely sometime before the transition from an aquatic to a land-based lifestyle, a sensory patch in this region of the inner ear was duplicated and subsequently adapted for sound detection. This adaptation has endowed vertebrates with a remarkably well tuned and sensitive ability to detect sound in air (Duncan and Fritzsch 2012).

The process of hearing via an inner ear, with some subtle variations, is fairly conserved in tetrapods. Pressure waves in the air are transmitted to the inner ear, which contains an organ known as the auditory papillae (cochlea in mammals) in which reside highly specialized hair cells that are sensitive to the resultant mechanical force (Gillespie and Müller 2009). Hair cells contain bundles of stereocilia at their apical end, which are stacked in a staggered fashion to respond to stimulation in a direction-dependent fashion (fig. 1C). Adjacent stereocilia are connected to one another by two members of the cadherin superfamily of cell-cell adhesion proteins, cadherin-23 (CDH23) and protocadherin-15 (PCDH15), which form a proteinaceous filament known as the tip link (Pickles et al. 1984; Ahmed et al. 2006; Kazmierczak et al. 2007) (fig. 1D). As with all members of the cadherin superfamily (Takeichi 1990; Kemler 1992; Shapiro and Weis 2009), CDH23 and PCDH15 are composed of a cytoplasmic domain, a transmembrane domain, and a variable number of extracellular cadherin (EC) “repeats”, with 27 in CDH23 and 11 in PCDH15 (fig. 1D). When stereocilia bend in response to sound or head movement, PCDH15 tugs at and applies tension in the cell membrane. This tension results in the transient opening of an ion channel and depolarization of the associated hair cell, followed by the transmission of an electrical impulse responsible for hearing and balance in a process known as mechanotransduction (Hudspeth and Corey 1977; Assad et al. 1991; Fettiplace and Kim 2014; Zhao and Müller 2015).

In mammals, it is well established that the tip link is composed of CDH23 and PCDH15, and that this interaction is required for mechanotransduction in the inner ear (Kazmierczak et al. 2007; Geng et al. 2013). This interaction is mediated through an interface between their two N-terminal EC repeats in a manner that is unique among cadherins (fig. 1D) (Sotomayor et al. 2012; Choudhary et al. 2020), and it has been validated by several biophysical experiments that have shown that the introduction of disease causing mutations breaks the complex (Sotomayor et al. 2012; Geng et al. 2013). The specificity of this interaction is such that a mutation in a single residue at the interface can abolish it completely, leading to a high level of conservation at such sites. Together, these results and biophysical experiments (Mulhall et al. 2021) revealed that the CDH23-PCDH15 interaction is strong enough to withstand the physiological forces that it would be subjected to in the inner ear. Furthermore, CDH23 and PCDH15 have been identified in zebrafish and chicken hair cells (Söllner et al. 2004; Seiler et al. 2005; Goodyear et al. 2010), suggesting that this interaction and its role in inner-ear mechanotransduction is conserved among vertebrates. While in the inner ear this complex exists as a heterotetramer in which a parallel dimer of CDH23 interacts with a parallel dimer of PCDH15 (Kazmierczak et al. 2007; De-la-Torre et al. 2018; Dionne et al. 2018; Choudhary et al. 2020), the locus of force transmission in this complex is the single CDH23-PCDH15 interaction formed between the N-terminal fragments (EC1-2) of the two proteins (subsequently referred to as the tip-link bond). As such, the single heterodimer composed of CDH23 EC1-2 and PCDH15 EC1-2, is an essential interaction for the transmission of mechanical force to the transduction channel complex of the associated stereocilia and is the focus of the present study.

Despite similarities in hair-cell mechanotransduction mechanisms mediated by tip links, there are several distinct variations in inner-ear anatomy between clades, such as papilla length, hair-cell arrangement and morphology, and innervation patterns (fig. 1A) (Coffin et al. 2004; Manley 2011). This diversity in anatomy suggests a likely diversity in the associated molecular machinery as well. However, little is known about the structure or mechanical properties of the tip link beyond that in mammals. Physiological and biochemical studies have been conducted on the tip link in chicken hair cells to characterize their functional elasticity in response to acoustic overstimulation (Husbands et al. 1999) and to identify key components and processes (Si et al. 2003; Ahmed et al. 2006; Goodyear et al. 2010; Indzhykulian et al. 2013), but there remains a dearth of biophysical or structural data. It is likely that this essential interaction has undergone mechanistic and structural modifications to accompany that of the associated auditory organs from fish to amniotes. Without further study of the CDH23-PCDH15 interaction and tip-link structure in these animals, it is difficult to propose a model of tip-link evolution in vertebrates. Because the auditory papillae in all extant lineages of vertebrates emerged and evolved independently in disparate biological contexts, the interface, structure, and strength of the tip link may vary significantly between clades.

While studying proteins in extant species provides a framework to guide evolutionary hypotheses, integrating information from the inferred ancestral state of biological molecules can greatly enrich such models and provide insight into protein function (Lynch et al. 2015; Zakas et al. 2017). One method used to probe the ancestral states of proteins is ancestral sequence reconstruction (ASR) (Pauling et al. 1963). Modern versions of this technique use maximum likelihood analysis, phylogenetic relationships, an alignment of extant protein or DNA sequences, and a model of sequence evolution to infer the most likely ancestral sequence that existed in the common ancestor of a single clade (for example, the ancestral amniote of fig. 1A) (Yang et al. 1995; Pupko et al. 2000). These ancestral sequences can then be used to produce the resurrected protein of interest and carry out experiments or to create models and run simulations, providing a glimpse of how it may have functioned in the evolutionary past of the lineage of interest and to observe evolutionary trends in protein function when compared to extant molecules (Lynch and Wagner 2008; Harms and Thornton 2013; Lim et al. 2018).

Because force plays such a prominent role in the function of the tip link, and the hearing abilities of vertebrates vary greatly between clades with a general increase in frequency sensitivity from fish to mammals, we initially hypothesized that tip-link bond strength would increase concurrently with the ability to integrate higher frequency sound, driven by sequence-level positive selection. To test this hypothesis, we used ASR, X-ray crystallography, experimental biophysical techniques, evolutionary sequence analyses, and molecular dynamics (MD) simulations to characterize the evolution of the tip-link bond in ancestral and non-mammalian species. While significant differences were found in tip-link bond strength, affinity, and dynamics between vertebrate species, there was limited class-wise correlation between tip-link strength and frequency sensitivity, contrary to the initial hypothesis. Instead, sequence level evolutionary changes, likely a result of neutral mutations that are constrained to varying extents by purifying selection and by inherent sequence-dependent properties of the proteins, manifests as lineage specific differences in interface dynamics, strength of interaction, and affinity. Results presented here allow the integration of molecular evolution and biophysics of the tip-link bond with that of the anatomical evolution of the inner ear and provide insight into the emergence of deafness-causing mutations in the tip link. Additionally, these results can be extended as a general model for the evolution of force-conveying protein complexes within and beyond the cadherin superfamily.

## RESULTS

### Ancestral Sequence Reconstruction

Sequences of the ancestral therian, ancestral eutherian, ancestral mammal (AncMam), and ancestral amniote (AncAm) CDH23 EC1-9 and the entire length of PCDH15 were inferred using ASR of amino acid sequences (see methods and supplementary dataset). The AncAm and AncMam CDH23 and PCDH15 EC1-2 were used in subsequent simulations and experiments (fig. 1E). The reconstructions of these ancestral states revealed several substitutions from the AncAm to the extant states that occurred at or near the interface that may have functional implications for the tip-link bond. From the AncAm to the AncMam, CDH23 Q138 was substituted for L at this position, which is found directly in the interface. From the AncMam state, L138 was retained in all subsequent animals in the mammalian lineage in the aligned species (fig. 1E). The reptiles, however, retained the ancestral Q138 at this position. Additionally, two tyrosines in CDH23 EC1 that are found directly in the interface, Y8 and Y11, were present in both the AncAm and AncMam, but were substituted for phenylalanine and histidine respectively in the mouse. Again, the reptiles retained the ancestral Y8 and Y11 states at these positions (fig. 1E), which further suggests that the reptilian and avian lineages have undergone fewer changes during evolution from the AncAm, and represent more ancestral states of the tip-link complex. The identity of the interface residues identified above were the same in the ancestral states regardless of the ASR method used (see methods).

### Evolutionary Divergence in Tip-Link Affinities

To determine if the sequence changes that occurred during evolution from the AncAm to extant vertebrates have functional effects, the lizard, mouse, and AncAm CDH23 and PCDH15 EC1-2 fragments were selected for experimental analysis using surface plasmon resonance (SPR). These complexes were chosen to cover a significant portion of vertebrate evolution, from the last common ancestor of the reptile and mammal lineage, represented by the AncAm complex, to the modern lizard and mouse. Using a kinetic analysis and a 1:1 binding model, the average *K_d_* values of the mouse, lizard, and AncAm were 3.6 ± 0.9 µM, 2.3 ± 1.0 µM, and 2.6 ± 0.1 µM, respectively (fig. 2A-C), indicating a slight but statistically significant decrease in affinity in the mouse complex relative to the other two (significance determined by estimation statistics and a 95% confidence interval of the mean difference between populations). The lizard exhibited a higher degree of non-specific interaction between the injected analyte (CDH23 EC1-2) and the SPR chip. This resulted in a smaller signal change after reference subtraction compared to the other two complexes, in addition to increased noise in the signal. However, this effect is below the detection limit of the machine, and the errors of the fits (a minimum of two orders of magnitude lower than the recorded value) were comparable to the other two complexes. As a negative control, I108N was introduced to the mouse and lizard PCDH15 EC1-2, as this mutation is known to cause deafness and break the tip-link interaction in mice (Geng et al. 2013). In both systems, this mutation completely abolished the response (supplementary fig. S1A and B), indicating that the interaction observed in the SPR experiments is the previously characterized handshake interaction (Sotomayor et al. 2012), and that this interface is conserved in non-mammals. Additionally, the on rate (*k_on_*) for the lizard complex was 3.0 ×10^4^ ± 4.3 M^-1^s^-1^, which was significantly higher than for the AncAm and mouse complexes (3.9 ×10^3^ ± 0.7 and 5.6 ×10^3^ ± 2.1 M^-1^s^-1^, respectively). This suggests that in equilibrium, evolution has modified the lizard interface to bind faster, which results in a higher affinity compared to the mouse complex.

**Fig. 2.**
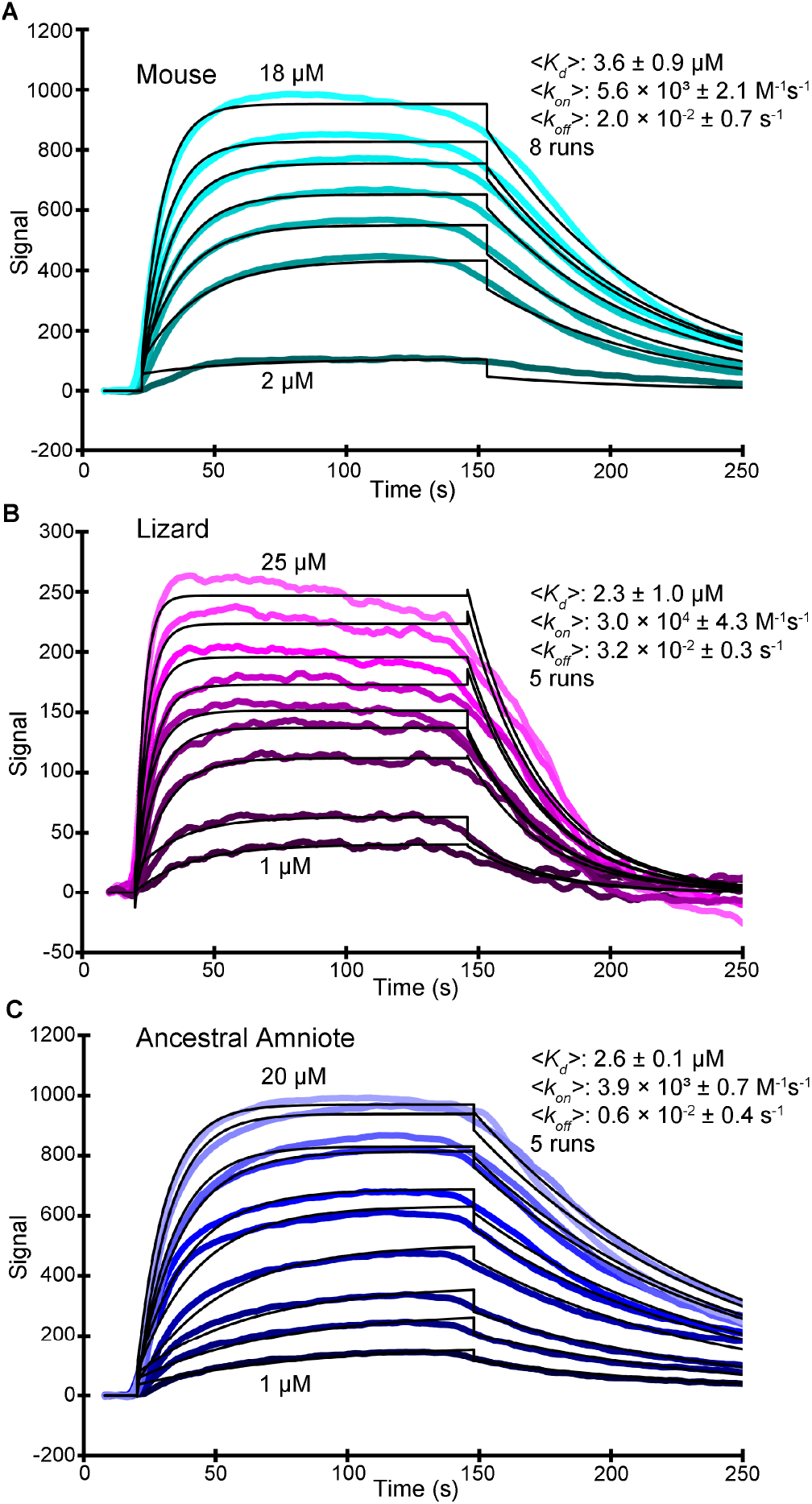
The mouse complex exhibits reduced affinity in SPR experiments. Representative curves from SPR experiments performed on the mouse (*A;* analyte injected at 2, 5, 8, 10, 12, 15, and 18 µM), lizard (*B;* analyte injected at 1, 2, 5, 8, 10, 12, 15, 18, and 25 µM), and AncAm (*C;* analyte injected at 1, 2, 3, 5, 8, 10, 12, 15, 18, and 20 µM) complex. Raw data are shown in the colored curves with the fits in black. The average affinities (*K_d_*), on rate (*k_on_*), and off rates (*k_off_*) are indicated, as well as the number of experiments performed on each complex. Values were obtained from a kinetic analysis of the raw data.

### Mammalian and Non-Mammalian CDH23 EC1-2 are Structurally Similar

To understand the structural effects of sequence variation in different vertebrate species we obtained four new crystal structures of CDH23 EC1-2, including those for the lizard (PDB ID 7N4P), AncAm (PDB ID 7SCM), AncMam (PDB ID 7SB6), and turtle (PDB ID 7SGX) (fig. 1F). We also included in our analyses a structure of the *Danio rerio* CDH23 EC1-2 (Jaiganesh et al. 2018). Sequence identities ranged from 73% to 97% suggesting that structures should be similar. To quantify structural similarity, we aligned the ancestral and non-mammalian structures to the crystal structure of the mouse CDH23 EC1-2 (PDB ID 2WHV) based on the C_α_ carbon atoms of residues 3 to 205 and computed root mean square deviations (RMSDs). The AncMam exhibited the lowest RMSD value compared to the mouse at 1.10 Å, followed by the AncAm at 1.26 Å, the turtle at 1.32 Å, and finally the lizard at 1.97 Å (supplementary fig. S2A, D, G, and J). While these results demonstrate some correlation between sequence and structural divergence, the low RMSD values for all structures aligned with the mouse indicate an exceedingly high structural conservation in CDH23 EC1-2 across vertebrate species, and throughout evolution from the ancestral amniote species to extant animals.

Next, the crystal structures of the lizard, AncAm, AncMam, and turtle CDH23 EC1-2 were aligned to the CDH23 EC1-2 monomer in the crystal structure of the mouse tip-link complex including PCDH15 EC1-2 (PDB ID 4APX). In all four structures, the aligned CDH23 EC1-2 structure fit in the tip-link interface without clashing with PCDH15 (supplementary fig. S2B, E, H, and K). All complexes formed by alignment of the crystal structure of CDH23 EC1-2 showed the interaction of a glutamine at position 98 of CDH23 (Q98) with an arginine at position 113 of PCDH15 (R113), an interaction that is known to be essential for the maintenance of this complex (Kazmierczak et al. 2007). Additionally, the lizard, AncAm, and turtle interface all exhibited a stabilizing CDH23 Q138-PCDH15 E111 interaction (explained further below) (supplementary fig. S2F, I, and L) while in the AncMam CDH23 Q138 was substituted to L138 during evolution, resulting in the loss of this stabilizing contact in the AncMam and mouse complexes (supplementary fig. S2C). The presence of these specific contacts, in addition to the high structural homology of the lizard, AncAm, AncMam, and turtle CDH23 EC1-2 crystal structures, suggest that the structure of the tip-link bond is conserved throughout the vertebrates, and that while certain essential interactions were maintained at the interface, evolutionary sequence changes have modified the network of interactions between CDH23 and PCDH15 EC1-2.

### Equilibrium All-Atom MD Simulations Reveal Species Specific Dynamics

Four systems were constructed and simulated using all-atom MD simulations, including a mouse (*Mus musculus*), lizard (*Anolis carolinensis*), pigeon (*Columba livia*) and AncAm CDH23-PCDH15 EC1-2 tip-link complex. These animals were chosen as representatives to cover a range of evolutionary divergence in vertebrates, from the ancestral state to mammals and reptiles. The mouse system was built directly from a crystal structure (PDB ID 4APX) (Sotomayor et al. 2012), while homology models of the lizard, pigeon, and AncAm were constructed from the mouse crystal structure (see methods). To determine how sequence diversity manifests in variation in structural and interface dynamics in this complex, each system was equilibrated for 83 ns (Sim1-4a, table 1).

**TABLE 1.** Overview of MD Simulations

| System | Label | Simulation time (ns) | Simulation Type | Speed (nm/ns) | Size (# atoms or # CG beads) | System Size (nm<sup>3</sup>) |
| --- | --- | --- | --- | --- | --- | --- |
| All-Atom Mouse | Sim1a-e | 471 | EQ/SMD | 0.1 | 202,310 | $28.6 \times 7.5 \times 8.7$ |
| All-Atom Lizard | Sim2a-e | 334 | EQ/SMD | 0.1 | 190,970 | $28.7 \times 8.1 \times 7.6$ |
| All-Atom AncAm | Sim3a-e | 396 | EQ/SMD | 0.1 | 209,058 | $28.9 \times 8.3 \times 8.3$ |
| All-Atom Pigeon | Sim4a-e | 298 | EQ/SMD | 0.1 | 171,602 | $27.9 \times 7.9 \times 7.4$ |
| CG Lizard | Sim5a-u | 20,661 | EQ/SMD | 0.1 | 14,278 | $25.9 \times 8.5 \times 8.1$ |
|  | Sim6a-p | 26,830 | SMD | 0.01 |  |  |
|  | Sim7a-e | 41,465 | SMD | 0.001 |  |  |
| CG Mouse | Sim8a-u | 15,911 | EQ/SMD | 0.1 | 14,278 | $25.9 \times 8.4 \times 8.0$ |
|  | Sim9a-p | 25,566 | SMD | 0.01 |  |  |
|  | Sim10a-e | 48,175 | SMD | 0.001 |  |  |
| CG Alligator | Sim11a-u | 3,292 | EQ/SMD | 0.1 | 14,269 | $25.9 \times 8.5 \times 8.0$ |
| CG AncAm | Sim12a-u | 3,360 | EQ/SMD | 0.1 | 14,270 | $25.9 \times 8.5 \times 8.0$ |
| CG Bearded Dragon | Sim13a-u | 3,173 | EQ/SMD | 0.1 | 14,283 | $25.9 \times 8.5 \times 8.7$ |
| CG Turtle | Sim14a-u | 2,855 | EQ/SMD | 0.1 | 14,275 | $25.9 \times 8.5 \times 8.1$ |
| CG Whale | Sim15a-u | 4,442 | EQ/SMD | 0.1 | 14,287 | $25.9 \times 8.5 \times 8.1$ |
| CG Bat | Sim16a-u | 3,622 | EQ/SMD | 0.1 | 14,280 | $25.9 \times 8.5 \times 8.9$ |
| CG Pigeon | Sim17a-u | 2,707 | EQ/SMD | 0.1 | 14,279 | $25.9 \times 8.5 \times 8.1$ |
| CG Fish1 | Sim18a-u | 2,622 | EQ/SMD | 0.1 | 14,266 | $25.0 \times 8.2 \times 7.8$ |
| CG Fish2 | Sim19a-u | 4,813 | EQ/SMD | 0.1 | 14,280 | $25.0 \times 8.2 \times 7.8$ |
| CG Fish3 | Sim20a-u | 4,266 | EQ/SMD | 0.1 | 14,273 | $25.0 \times 8.2 \times 7.8$ |
| All-Atom Mouse | Sim21 | 50.8 | Constant Force | - | 202,310 | $28.6 \times 7.5 \times 8.7$ |
| All-Atom Lizard | Sim22 | 58.3 | Constant Force | - | 190,970 | $28.7 \times 8.1 \times 7.6$ |
| All-Atom AncAm | Sim23 | 53.4 | Constant Force | - | 209,058 | $28.9 \times 8.3 \times 8.3$ |

The buried surface area (BSA) between CDH23 and PCDH15 was measured during equilibration for all simulations. BSA values have implications for binding affinity in a variety of protein complexes, where in general an increase in affinity is observed with increases in BSA values between components of a complex (Chen et al. 2013). In the first 20 ns of the equilibration, the lizard, AncAm, and pigeon exhibit an increase in BSA, from ∼1,125 Å^2^ in each to ∼1,250 Å^2^. However, no increase in buried surface area is observed in the mouse complex during this time as it remains at ∼1,125 Å^2^ (fig. 3A). Lineage specific differences in sequence are responsible for this difference in structural dynamics. In AncAm, lizard, and pigeon, CDH23 Y7 interacts with PCDH15 R216 while CDH23 Y11 interacts with PCDH15 R117. This facilitates a local motion in which the N-terminus of CDH23 closes on EC2 of PCDH15, resulting in the increase in buried surface area in these two species (fig. 3A-C) and stable interactions throughout the equilibration (supplementary fig. S3A, C and D). The mouse CDH23 sequence instead has a phenylalanine and a histidine residue at positions 7 and 11, respectively (fig. 1 E), and so this complex did not form the favorable interactions seen in the other three complexes. As a result, the distance of the N-terminus in CDH23 did not change appreciably relative to EC2 of PCDH15 in the mouse complex (fig. 3C), which precluded stability in these interactions during the entire equilibration (supplementary fig. S3B). After 20 ns, the surface area of the mouse increases to a value similar to that of AncAm and the lizard. However, unlike the AncAm, lizard, and pigeon, this increase is due to a fluctuation in the N-terminus of PCDH15, which forms a closer association with EC2 of CDH23 that is not observed in the AncAm, lizard, or pigeon. The results from the equilibrium simulations demonstrate that sequence-level changes can result in unique interface dynamics, with the lizard and pigeon exhibiting a closure of CDH23 EC1 on PCDH15 EC2 similar to AncAm, while the mouse did not. This suggests that a closer association between CDH23 and PCDH15 in this region may represent an ancestral trait for this complex.

**Fig. 3.**
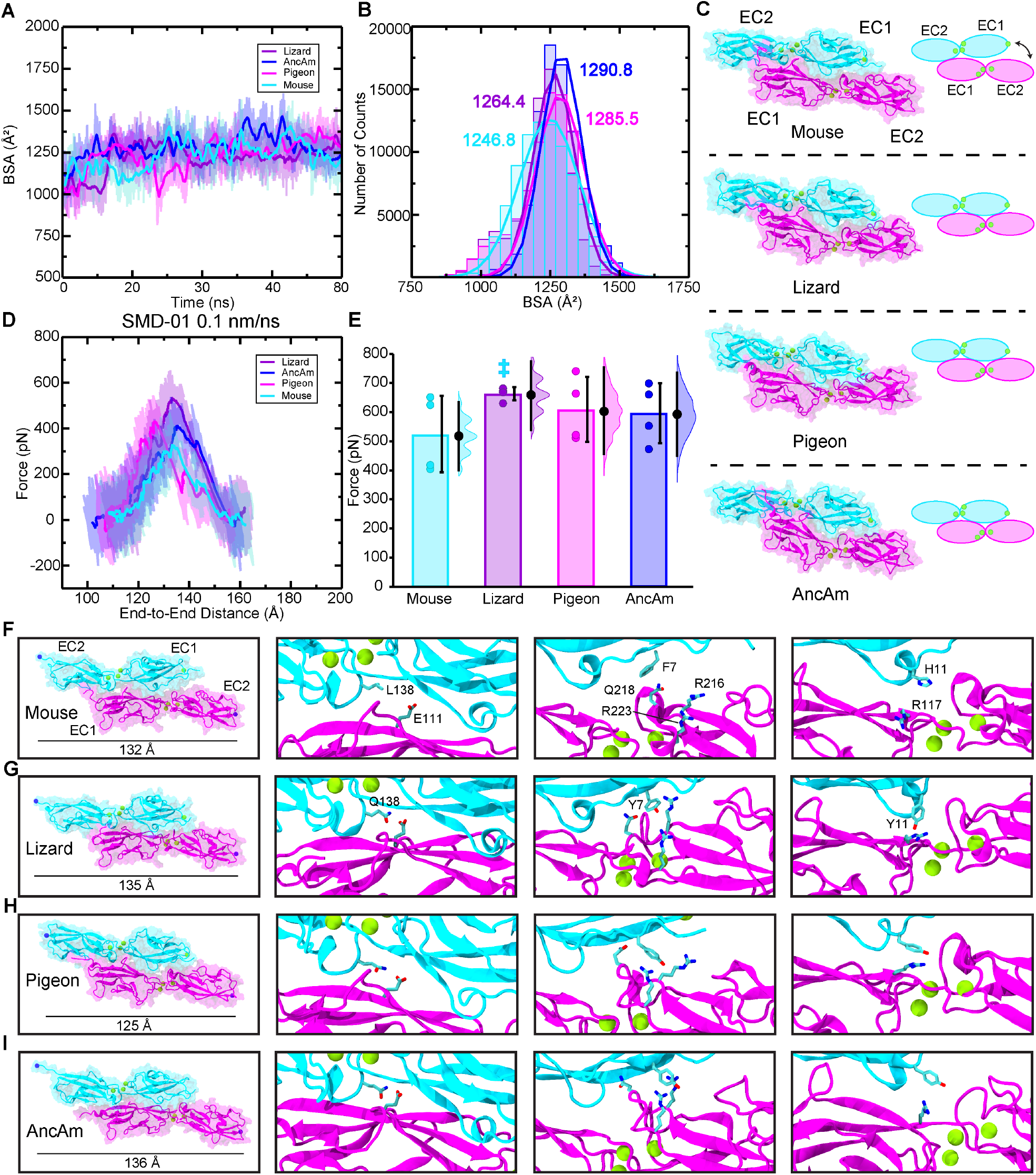
All-atom MD simulations reveal lineage specific differences in tip-link bond dynamics and strength due to sequence changes. (*A*) BSA is shown for all four complexes during 83 ns long equilibrations (Sim1a, Sim2a, Sim3a, and Sim4a). (*B*) Count distributions from (*A*) with associated fits to the normal distribution. Mean values are indicated with the respective colors. (*C*) Structures of each complex at the end of the 83 ns are shown in ribbon representation, with their molecular surfaces in transparent representation. Ca^2+^ ions are shown as green spheres. Schematics to the right show how the orientation of mouse CDH23 EC1 and PCDH15 EC2 remained more separated relative to the other three species. (*D*) The force extension profile for each complex for the first SMD simulation (Sim1b, Sim2b, Sim3b, and Sim4b). The mouse exhibited the lowest force peak, followed by the AncAm and Pigeon, and the lizard exhibited the highest force. (*E*) Individual force peak values, averaged between both pull atoms, and distribution for all four SMD simulations are shown, as well as their average by the height of the box. Standard deviations are shown by the single black lines, while the 95% confidence interval from estimation statistics are shown by the black lines and circles to the right of each bar, with the resampling distribution of the differences in means on the right. The lizard exhibited a significantly higher force than the mouse (as measured by estimation statistics) indicated by the cyan dagger. (*F-I*) On the left is the structure at the force peak of the first SMD simulation for each complex. The next three panels indicate the three interactions in the interface that differ between reptiles and mammals, and that contribute to differences in interface dynamics. Residues in PCDH15 are identical for all four complexes and are labeled in (*F*). Residues in CDH23 that differ from the mouse are indicated for the lizard in (*G*) and are identical for the pigeon and AncAm.

Next, equilibrium simulations of all three systems were analyzed using network analysis (Vendruscolo et al. 2001; Sethi et al. 2009; Krieger et al. 2020). Combined with MD simulations, network analysis was recently used to identify residues that disrupt the path of force transduction in the mammalian tip-link (Hazra et al. 2019). In these models of protein structure, each residue in the protein represents a node and connections between nodes define the overall topology of the physical network. Here, network analysis was used to determine if the connectivity between and within CDH23 and PCDH15 is affected by the sequence divergence observed between vertebrate lineages. Computed over the 83 ns of equilibration, the network topology varied between the three species. In particular, a CDH23 Y7 residue forms interactions with residues on PCDH15 in the lizard, pigeon, and AncAm systems (fig. 3F-I, supplementary fig. S4D, G, and J). As noted above, the mouse experienced a Y7F substitution during evolution, and this resulted in a reduced connectivity between CDH23 and PCDH15 as revealed by network analysis (supplementary fig. S4A, D, G, and J), in agreement with the observed lower buried surface area of the mouse complex in equilibrium (fig. 3A and B). Next, the optimal path from the C-terminus of CDH23 to the C-terminus of PCDH15 was calculated during equilibration. For all four complexes, the optimal path between C-termini is nearly identical, and all residues that lie along this path are mostly conserved, with minor exceptions at residues away from the interface (supplementary fig. S4B-C, E-F, H-I, and K-L). This high conservation suggests that this path of communication is important for force transduction in the tip link and has not been altered during evolution, as the residues that comprise this path are maintained. Instead, substitutions in the surrounding interface have been utilized by evolution to modulate tip-link strength and dynamics without causing detrimental changes to the path of force propagation.

### Steered MD Shows Significant Changes in Tip-Link Strength as a Result of Sequence Divergence

Force was applied to the CDH23-PCDH15 complex in four systems using SMD to predict how sequence evolution affects interaction strength and unbinding interface dynamics of the tip link. We ran four constant-velocity SMD simulations at 0.1 nm/ns for each system (Sim1-4b-e, table 1) starting from coordinates saved at different timepoints in the corresponding equilibrium simulations.

The average forces and distribution of individual force-peak values for each complex in all four simulations is shown in fig. 3E. The average force for the lizard complex (*F*_p_ = 667.6 ± 22.8 pN) was significantly higher than the mouse complex (*F*_p_ = 527.5 ± 132.4 pN) analyzed with estimation statistics, but not with a t-test (*P* = 0.05). The average force peaks for the AncAm complex (*F*_p_ = 598.8 ± 103.4 pN) and pigeon complex (*F*_p_ = 611.8 ± 112.8 pN) were both higher than the mouse, but the results were not statistically significant. While significance is difficult to obtain with the limited sampling achieved with all-atom MD, these results suggest that lineage-specific sequence divergence affects protein function not only in equilibrium, but also in a force-dependent context, with the lizard complex requiring the highest force to break, and the mouse complex requiring the lowest force.

To gain further insights into the molecular determinants of the variable tip-link strength between species, we analyzed SMD trajectories from four repeat simulations individually. The force-extensions profiles from the first SMD simulation are shown in fig 3D. For clarity, only the force experienced at the C-terminal C_α_ carbon of CDH23 is shown. Each was started from coordinates saved at 21 ns in the equilibrium simulation. The lizard complex exhibited the highest force peak at 634.5 ± 6.2 pN, followed by the Pigeon at 513.1 ± 33.1 pN, the AncAm at 474.8 ± 8.3 pN, and finally the mouse at 397.9 ± 11.8 pN, averaged between the force experienced at the C-terminal C_α_ carbons of CDH23 and PCDH15 in each case. As seen in fig. 3A, the buried surface area at 21 ns of the lizard, pigeon, and AncAm complex is ∼ 1,250 Å^2^, while that of the mouse is ∼ 1,125 Å^2^. Additionally, during the entire 83 ns equilibration, the mouse exhibited the lowest average BSA (fig. 3B). This difference in interfacial BSA, which was shown to result from lineage specific sequence differences (fig. 1E), may contribute to the force response observed in each complex. In addition to the BSA increase, several interactions unique to the AncAm and reptile lineage persist until the force peak in this simulation. In lizard, pigeon, and AncAm, CDH23 Q138 interacts with PCDH15 E111, CDH23 Y11 with PCDH15 R117, and CDH23 Y7 with PCDH15 Q218. In the mouse, CDH23 Q138L, Y11H, and Y7F substitutions destabilize these three interactions (fig. 3F-I, supplementary figs. S5-S8A-C). While these three interactions appear to be stable at the force peak for the lizard complex, and to a lesser extent for the AncAm and pigeon complexes, another set of interactions unique to non-mammals only forms closer to the force peak. An asparagine at position 2 of PCDH15 in lizard and AncAm was substituted for a tyrosine in the mammalian lineage, and upon the application of force this asparagine forms transient interactions with CDH23 in AncAm, lizard, and pigeon. An additional interaction was observed after unbinding in the lizard complex between CDH23 R4 and PCDH15 E111 that correlated with an increase in force after the force peak. Overall, these lineage-specific interactions alter the response to force in the tip link and correlate with higher forces of unbinding in non-mammals.

The second SMD simulation repeat for each complex was started from coordinates saved at 23 ns in the equilibrium simulation, except for the pigeon which was started at 72 ns, chosen arbitrarily. The AncAm and pigeon complexes exhibited increases in force peak values at 555.0 ± 32.5 pN and 666.5 ± 16.9 pN, respectively. Unlike the first simulation, the lizard and mouse exhibited similar force peaks at 677.8 ± 24.5 pN and 655.0 ± 9.4 pN, respectively. The mouse complex experienced a force-dependent increase in BSA during the first 10 ns of the simulation (supplementary fig. S5E). This force-dependent increase in BSA results from a rolling motion in which PCDH15 becomes more closely associated with CDH23 between EC1 and EC2, and likely accounts for the comparable force peak between lizard and mouse. This behavior is characteristic of a catch-bond, and these results suggest that the tip link is capable of this response.

For the starting point of the third SMD simulation repeat, starting coordinates were arbitrarily taken at 66 ns, 72 ns, 39 ns, and 69 ns for the AncAm, lizard, pigeon, and mouse equilibrium simulations, respectively. The AncAm, pigeon, and lizard exhibited increases in force peak values compared to the previous two simulations at 663.6 ± 2.7 pN, 744.3 ± 19.8 pN, and 686.0 ± 35.9 pN, respectively. The Mouse exhibited a force peak of 420.3 ± 33.0 pN, similar to the first simulation. The fourth and final SMD simulation repeat was started using the same starting coordinates as the third SMD simulation repeat for the AncAm, lizard, and pigeon complexes, while the mouse was started from coordinates saved at 64 ns of its equilibrium simulation. The force peaks for the AncAm, lizard, mouse, and pigeon were 701.7 ± 13.5 pN, 672.2 ± 33.2 pN, 628.3 ± 36.5 pN, and 523.1 ± 18.0 pN, respectively. While this variability demonstrates that the force response is sensitive to the starting configuration, the average difference between the lizard (*F*_p_ = 667.6 ± 22.8 pN) and mouse (*F*_p_ = 527.5 ± 132.4 pN) remained significant analyzed with estimation statistics.

To further study the mechanical properties of the tip-link bond, we also carried out constant force pulling simulations. Upon application of a 150 pN constant force, the lizard and AncAm complex exhibited a higher stiffness compared to the mouse (313.8, 127.1, and 90.3 pN/nm, respectively). The lizard and AncAm also exhibited a higher damping coefficient compared to the mouse (113.1, 226.2, and 62.4 pN ns nm^-1^). These results suggest that a less elastic interface that is more resistant to mechanical perturbations may be an ancestral state of the tip-link complex (supplementary fig. S9A). In addition to differences in the interface highlighted above for constant-velocity pullings, analysis of the constant-force trajectories revealed variation between species that may contribute to this difference in force response. In the lizard and AncAm, there is a direct intermolecular interaction between CDH23 D14 and PCDH15 R117. In the mouse however, a CDH23 G93R substitution results in an intramolecular interaction between D14 and R93, preventing it from forming an intermolecular contact. Additionally, the reduced contact between CDH23 EC1 and PCDH15 allows a hydrogen bond to form between the CDH23 P6 backbone oxygen and the PCDH15 R216 sidechain in the AncAm and lizard, but not in the mouse (supplementary fig. S9B). Such differences may contribute to the less elastic force response exhibited by the AncAm and lizard complexes.

### Exploring Tip-Link Strength in an Expanded Subset of Species Over Longer Timescales With Coarse-Grained Simulations

To reach the microsecond time scale relevant for auditory transduction and to carry out multiple repetitions of SMD simulations for an expanded subset of vertebrate species, we used coarse-grained (CG) MD (Joshi and Deshmukh 2021). First, each system was minimized and equilibrated for 23.6 ns. From this equilibrated structure, each system was pulled at 0.1 nm/ns using SMD in 10 independent simulations. The equilibration was extended to 283 ns for each complex, and an additional 10 pulling simulations at 0.1 nm/ns were carried out from different starting configurations along the equilibration, for a total of 20 SMD simulations per system (∼20,000 ns total per system). The lizard and mouse complexes were additionally pulled at 0.01 nm/ns in 16 independent runs (∼25,000 ns total per system), 7 of which began from the same starting coordinates as the first 10 0.1 nm/ns pullings and the remining 9 were started from the extended equilibrium simulations. Finally, the mouse and lizard complexes were pulled at 0.001 nm/ns in five independent runs adding up to ∼45,000 ns for each system, all starting from different points along the equilibration.

The results of the 0.1 nm/ns pulling simulations are shown in fig. 4A. Force required to unbind these complexes showed significant variability within and between lineages. As with the all-atom simulations, sequence changes in the interface are responsible for this variability in complex strength. This is seen in fig. 4D-G, where the interface of the whale, lizard, yellowtail fish, and turtle are shown in detail, respectively, at the force peak. Highlighted in orange are the residues that differ from that of the mouse and may contribute to the significant difference in force exhibited by these complexes.

**Fig. 4.**
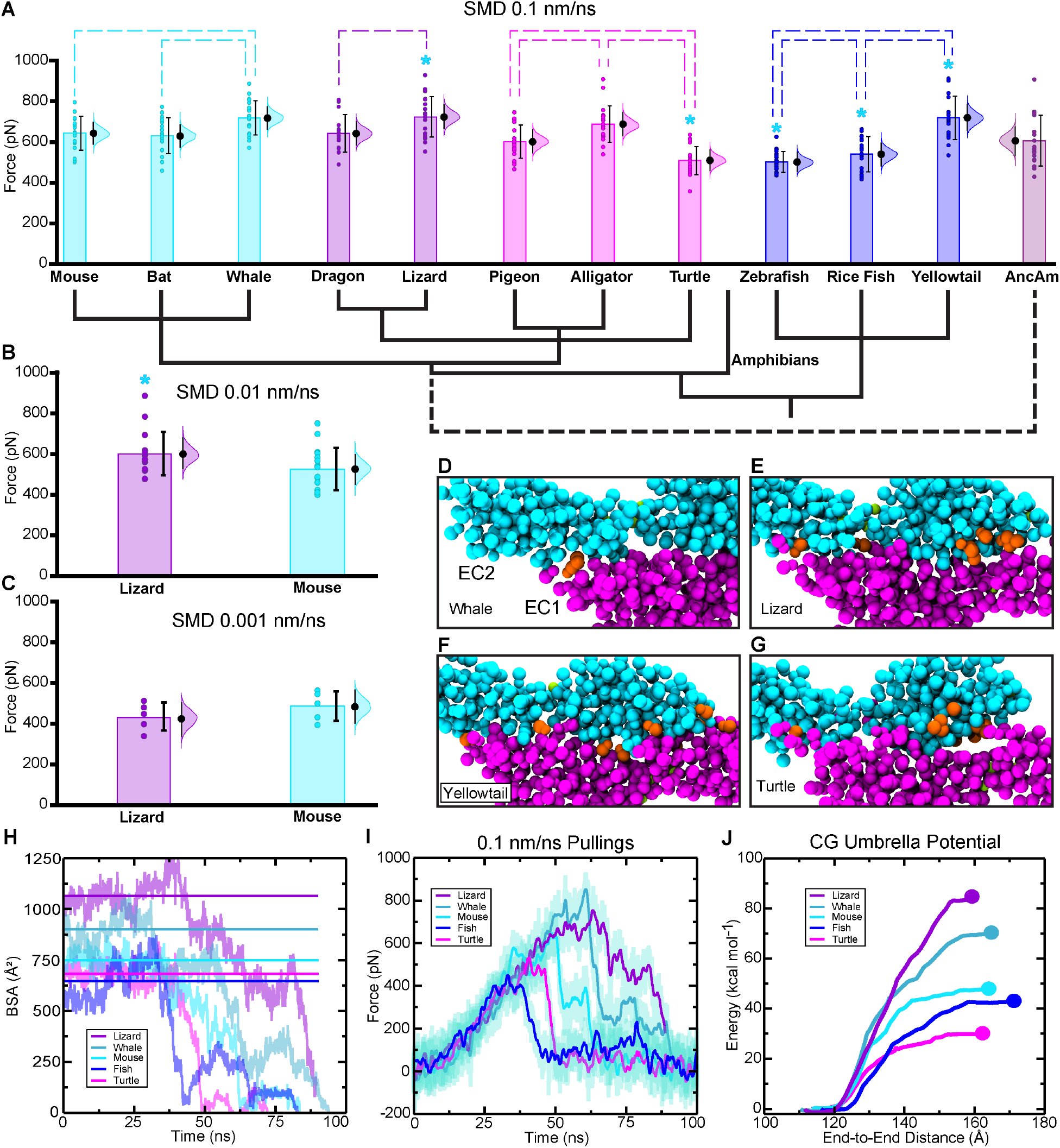
CG MD simulations show trends in tip-link properties across vertebrates that correlate with BSA and interface energy. (*A*) Results of CG SMD at 0.1 nm/ns (Sim5b-u, Sim8b-u, Sim11-20b-u). Data points are shown in circles and distributions are shown for each complex as in fig. 3 (*N* = 20), as well as the average indicated by the height of the box. Significant differences (determined from t-test and estimation statistics) from the mouse are indicated by a cyan star, while significant differences within lineages are indicated by dashed lines. (*B*) Results of CG SMD simulations at 0.01 nm/ns for the lizard and mouse (Sim6a-p and Sim9a-p) shown as in (*A*). The lizard exhibited a significantly higher average force peak. (*C*) Results of CG SMD simulations at 0.001 nm/ns for the lizard and mouse (Sim7a-e and Sim10a-e) shown as in (*A*). At this pulling speed, the lizard and mouse did not exhibit significantly different average force peaks. (*D-G*) Each complex shown exhibited a significantly different average force peak from the mouse at 0.1 nm/ns, and sequence differences at the interface from the mouse are indicated in orange. (*H*) The BSA for the first SMD simulation of each complex indicated is shown as raw data (curves) and as the average BSA before the drop upon unbinding (solid horizontal line). (*I*) Force profile for the same simulations as in (*H*). (*J*) The energy required to unbind each complex, calculated as a potential of mean force from umbrella sampling CG simulations. The same simulations were used to analyze the BSA, the force for each complex, and to generate initial conformation to compute the energy of unbinding.

In agreement with the all-atom SMD simulations, the lizard had a significantly higher average force peak at 0.1 nm/ns (722.8 ± 99.4 pN) than the mouse (641.6 ± 84.4 pN), while the mouse and AncAm (605.4 ± 123.8 pN) were not significantly different (*P* = 0.01 and *P* = 0.17, respectively). Similarly, at 0.01 nm/ns, the lizard had a significantly higher average force peak (603.1 ± 107.2 pN) than the mouse (526.6 1 ± 105.0 pN; *P* = 0.02), though the difference is less than at 0.1 nm/ns (fig. 4B). Finally, at 0.001 nm/ns, the lizard and mouse force peaks were not significantly different at 440.4 ± 69.0 pN and 492.5 ± 73.4 pN, respectively (fig. 4C; *P* = 0.09). This varying response at different speeds may reflect varying degrees of purifying selection between the mouse and lizard, as the lizard inner ear is not physiologically capable of integrating high frequency sound. Because of this, the lizard tip link would not be under selection to break in response to high frequency, high amplitude sound, while the mouse may have undergone purifying selection for the tip link to break before damage is incurred to the protein or surrounding tissue.

As with the all-atom simulations, BSA was analyzed in a subset of the CG-SMD simulations to determine if it could correlate with the significant differences in force peaks. A representative subset of the simulated systems was selected for further analysis: the turtle, mouse, zebrafish, lizard, and whale. BSA was analyzed throughout one SMD simulation at 0.1 nm/ns in each of these animals (fig. 4H). The average buried surface area of the initial phase, before the first permanent decrease due to unbinding, is shown in the solid horizontal line for each animal. With minor exceptions, this value correlates with the force required to unbind the complex in each animal for the associated SMD simulation (fig. 4I), with higher BSA correlating with higher force peaks. The lizard exhibited the highest buried surface area (1,066.1 Å^2^) and second highest force peak (740.0 ± 21.9 pN, averaged between force from both C-terminal backbone particles), followed by the whale (901.4 Å^2^, 837.9 ± 23.0 pN). While the force peak was higher for the whale in this simulation, the average of all 20 simulations for the lizard and whale are not significantly different (722.8 ± 99.4 pN and 719.0 ± 83.5 pN respectively). The mouse exhibited intermediate values for both buried surface area (749.0 Å^2^) and force (645.3 ± 2.4 pN), followed by the turtle (682.3 Å^2^, 535.5 ± 5.5 pN) and zebrafish (646.6 Å^2^, 447.2 ± 4.1 pN). This correlation between force peak and buried surface area in each system suggests that, as with the all-atom simulations, lineage specific differences in protein sequence affect interface dynamics which in turn affects the force required to unbind the tip-link, and that more evolutionarily primitive animals with simpler inner-ear structures (zebrafish and turtle) exhibit the lowest forces required to unbind the tip-link complex, likely as a result of their lowered BSA.

### Species-Specific Energies of Unbinding Correlate With Tip-Link Strength and BSA

The energy of unbinding for the tip-link bond was measured using umbrella sampling in the five systems for which BSA was analyzed (turtle, mouse, zebrafish, lizard, and whale). In this technique, configurations are generated along a certain reaction coordinate (in this case the end-to-end distance for the complex), and independent constrained simulations are ran starting from each of those configurations. From these independent simulations, an energy profile can be reconstructed, which in this case provides the energy required to unbind the complex (You et al. 2019). Results for each of the five systems are shown in fig. 4J. The lizard exhibited the highest energy of unbinding (85.40 kcal mol^-1^; 3,211 ns of sampling over 32 windows), followed by the whale (71.04 kcal mol^-1^; 3,311 ns of sampling over 33 windows), mouse (47.97 kcal mol^-1^; 3,311 ns of sampling over 33 windows), zebrafish (43.49 kcal mol^-1^; 3,411 ns of sampling over 34 windows), and turtle (30.16 kcal mol^-1^; 3,110 ns of sampling over 32 windows). While the CG model is limited in its ability to capture accurate free energies and these are not values expected for experimental or *in vivo* energies for the tip-link interaction, they can be used to establish a hierarchy of interaction strengths. As with force of unbinding, the energy required to unbind each complex correlates with BSA, except for the zebrafish and turtle, which are also not significantly different averaged over the 20 pulling simulations. This further supports the assertion that BSA is correlated with the observed variety in tip-link strength, and as with force the two most basal animals (zebrafish and turtle) required the lowest energies to unbind the complex.

### Evolutionary Sequence Analyses Reveal How Biophysical Properties Influence Rate of Evolution

While studying extant protein homologs can explain how evolutionary sequence changes in a protein affect its function, this approach alone is limited in its ability to predict how biophysical properties shape the underlying molecular evolution in different genetic contexts. Molecular evolution simulations (Kohn et al. 2005; McCandlish and Stoltzfus 2014; Nelson and Grishin 2014) that incorporate protein structure can determine how the biophysical properties of CDH23 and PCDH15 constrain the evolutionary sequence landscape in different homologs, and can provide an evolutionary explanation for the results observed in the MD simulations and experiments (Xia and Levitt 2004; Teufel et al. 2019). Evolutionary trajectories were simulated for the CDH23-PCDH15 EC1-2 complex and its path through the fitness landscape was traced by evaluating the fitness of each mutation using FoldX energies (Schymkowitz et al. 2005) at each step and by selecting or rejecting each mutation with a probability function modified by its fitness. Such simulations have been used to study sequence and structural divergence in proteins from distantly related species (Kachroo et al. 2015). This approach has been used here to reveal how biophysical properties can constrain the rate of molecular evolution, providing a link between the physical and evolutionary properties of cadherins.

In the evolutionary simulations, a monomorphic population is assumed (Teufel and Wilke 2017) and the number of mutations that were fixed (accepted and inserted in the structure) per site was taken as the evolutionary rate for each protein of each species. Simulations were analyzed on two timescales; at 3000 accepted mutations, corresponding to a long-term steady state (long timescale), and at 75 accepted mutations, corresponding roughly to the amount of time since the last vertebrate ancestor (∼ 500 million years; vertebrate timescale), determined based on the number of substitutions between the AncAm and extant mouse CDH23 and PCDH15 EC1-2 sequences, which are separated by ∼ 300 million years. If a subset of residues in the complex had accumulated more mutations than another subset throughout the 3000 or 75 steps of each run, it can be assumed that they are more robust to mutations and would accommodate mutations more readily during evolution. Mutations were first divided into two classes; those at the interface, defined as any residue that is within 6 Å of its binding partner in the starting structure, and non-interface sites, defined as any residue not at the interface. Number of accumulated mutations was then normalized by dividing by the total number of sites in the interface for interface substitutions, or by the number of non-interface sites for non-interface substitutions to give average number of mutations per residue. Because calcium (Ca^2+^)-binding residues are completely conserved in all species considered and are required for cadherin function, these residues were excluded from all analyses.

Contact density (CD) was monitored during MD and evolutionary simulations, as it has been shown to correlate with evolutionary rates in proteins (Bloom et al. 2006; Zhou et al. 2008). Higher CD will increase the stability of globular proteins, making them more robust to mutations. CD has also been shown to be a determinant of sequency entropy (Shakhnovich et al. 2005), which measures the number of sequences that adopt a particular structure. Supplementary table S1 shows the normalized number of accumulated non-interface mutations over each trajectory for each protein fragment (either CDH23 or PCDH15 EC1-2), the CD for that protein during equilibrium MD, and the CD for that protein during the evolutionary simulation at both timescales. The average CD was 17.25 ± 0.99 for CDH23 EC1-2 and 13.97 ± 0.36 for PCDH15 EC1-2 in equilibrium MD, and 14.72 ± 0.23 for CDH23 EC1-2 and 14.27 ± 0.08 for PCDH15 EC1-2 in evolutionary simulations over 3000 steps (averaged across species, supplementary fig. S10A-H), while the average number of fixed non-interface mutations during evolutionary simulations was 7.37 ± 0.02 for CDH23 EC1-2 and 6.22 ± 0.01 for PCDH15 EC1-2 over 3000 steps (supplementary table S1). These differences are all significant (*P* < 0.05), which suggests that more compact (i.e., more stable or high sequence entropy) structures are more robust to mutations and will evolve at faster rates due to different structural constraints. The average CD was 14.55 ± 0.18 for CDH23 EC1-2 and 14.21 ± 0.11 for PCDH15 in evolutionary simulations over 75 steps, while average number of fixed non-interface mutations was 0.247 ± 0.089 for CDH23 EC1-2 and 0.232 ± 0.078 for PCDH15 EC1-2 over the same timescale (supplementary table S1), demonstrating a correlation between CD and evolutionary rate at timescales relevant for vertebrate evolution. This effect is apparent regardless of location in the protein, with CDH23 experiencing an increase in fixed mutations in both interface and buried sites in all species (supplementary fig. S11) and is maintained both during the timescale that is relevant for protein function as revealed by MD simulations (supplementary fig. S10E-H), and over evolutionary timescales suggested by the evolutionary simulations (supplementary fig. S10A-D). Interestingly, during the vertebrate timescale simulation, the difference in average number of fixed mutations between CDH23 and PCDH15 was greatest in the AncAm (supplementary table S1), suggesting that this rate difference was prominent during the transition from the ancestral to extant amniotes.

Results from evolutionary simulations and CD analyses also agree with evolutionary rate analyses, which showed an average rate of 1.22 for CDH23 EC1-2 and 1.11 for PCDH15 EC1-2 considering all sites using an amino-acid based model and the LEISR method (Spielman and Pond 2018). Considering only buried sites, which will generally represent the slowest evolving residues in globular proteins (Bloom et al. 2006), the average rate was 0.55 for CDH23 EC1-2 and 0.29 for PCDH15 EC1-2. A higher rate of evolution for CDH23 EC1-2 is also indicated by analyzing the sequence identity and similarity between species, with a mean identity of 81.7 % ± 8.2 for CDH23 EC1-2 and 88.0 % ± 8.4 for PCDH15 EC1-2 and a mean similarity of 82.1 % ± 8.5 for CDH23 EC1-2 and 91.6 % ± 6.4 for PCDH15 EC1-2, using the same alignments as in the evolutionary rate analyses. Finally, a BUSTED analysis reveals a significant difference in rate distribution between CDH23 EC1-2 and PCDH15 EC1-2 (likelihood ratio test (LRT) = 24.7, *P* = 0.00016), with a greater proportion of CDH23 EC1-2 (0.1%) under strong positive selection than PCDH15 EC1-2 (0.01%). Additionally, a greater proportion of PCDH15 EC1-2 (10.2%) is under strong purifying selection than CDH23 EC1-2 (8.2%). These results strongly suggest that the rate of evolution of cadherins is constrained by the compactness of the EC repeats from which it is composed, as demonstrated by the correlation between evolutionary rate and CD in CDH23 EC1-2 and PCDH15 EC1-2. This increased rate of evolution in CDH23 EC1-2 further suggests a greater propensity for evolutionary innovation in CDH23 relative to PCDH15, which is reflected in the substitutions at the interface of the mouse complex relative to the ancestral state that might be responsible for the divergent properties observed in MD simulations (fig. 3, fig. 4) and all of which are present in CDH23. Finally, the apparent higher rate of evolution in CDH23 EC1-2 may suggest an underlying biophysical explanation for the observation that CDH23 EC1-2 is host to far more deafness causing mutations than PCDH15 EC1-2 (Jaiganesh et al. 2017). The relaxed biophysical constraints on CDH23 EC1-2 implied in the evolutionary simulations here may permit such fitness-lowering mutations, and this is particularly evident when comparing the fixed mutations distributed by residue between CDH23 and PCDH15 during the evolutionary simulations (supplementary fig. S11).

The average number of accumulated mutations for the interfaces of each species and the average interface contact density during the MD equilibration for each species (supplementary table S2) demonstrate an inverse correlation. The lizard, AncAm, and pigeon exhibited significantly fewer fixed mutations per residue in the interface during the evolutionary simulation evaluated over 3000 steps (6.97, 7.24, and 7.25 respectively) compared to the mouse (7.32). Concurrently, the lizard, AncAm, and pigeon exhibited higher interface CD in equilibrium MD (11.67, 11.70, and 11.79 respectively) compared to the mouse (11.42), as well as during the evolutionary simulation over 3000 steps (supplementary fig. S10I-L). Over 75 evolutionary steps, the lizard, AncAm, and pigeon again exhibited the highest interface CD (11.71, 11.53, and 11.57 respectively) relative to the mouse (11.44), as well as fewer fixed interface mutations per residue (0.155, 0.178, and 0.178 for the lizard, AncAm, and pigeon, respectively) compared to the mouse (0.184; supplementary table S2). This implies that a more compact or crowded interface is less able to accommodate mutations during evolution. This effect has been previously reported, where the degree of interface burial upon complex formation was shown to correlate with a decrease in evolutionary rate (Eames and Kortemme 2007; Franzosa and Xia 2009). Because this property was apparently decreased during mammalian evolution, the mouse complex may have been more susceptible to substitutions at the interface relative to the ancestral and reptilian complex, leading to the divergent properties between the mouse and lizard complexes observed in MD simulations and experiments.

The propensity of certain amino acids to be fixed in these simulations varied between interface and non-interface residues, between CDH23 and PCDH15, and between species (supplementary fig. S11). The lizard, pigeon, and AncAm all exhibited increases in fixed mutations in CDH23 buried sites relative to the mouse (supplementary fig. S11A, E, I, and M), and also exhibited higher overall CD than the mouse both in evolutionary simulations and MD simulations (supplementary fig. S10A-H). This increase in overall CD may allow CDH23 in these species to accommodate the loss of stability due to the insertion of bulkier or polar residues such as ARG, which was fixed more often in all three non-mouse species, or GLU, which was fixed more often in the lizard and AncAm. Additionally, there was some evidence for reduction in rate of fixation in interface sites in the non-mammalian species, all of which exhibited greater interface CD in evolutionary and MD simulations (supplementary fig. S10I-L). This supports the hypothesis that a more crowded interface would be less likely to accommodate mutations, which is also indicated by the inverse correlation between interface CD and interface residue evolutionary rate.

There was a difference in the distribution of residues accepted at interface vs. buried residues during the evolutionary simulations. For all species, a large relative decrease in the acceptance of large or polar amino acids is observed in buried sites compared to interface sites (supplementary fig. S11 *A, C, E, G, I, K, M,* and *O*), compared to hydrophobic residues. Insertion of residues such as ARG, LYS, and GLU near the interface usually resulted in an increase in BSA, whether the energy of the interface increased or decreased. This implies that even when it is not explicitly selected for, BSA of an interface and by extension its mechanical strength will increase during evolution in the absence of explicit selection against it, providing a possible explanation for how BSA and higher unbinding strength can emerge from neutral evolution.

The previous analyses were performed only on variants that were fixed during the simulation, to represent the evolutionary trajectory of the genotype that would be dominant in a monomorphic population. The evolutionary simulations were also ran keeping track of each attempted mutation for 312 total attempted mutations, with subsequent analysis performed on structures corresponding to the fixed variant but weighted by the number of steps in which that variant was present as the dominant genotype. The results of the simulations analyzed in this way agree with all previous analyses, with CDH23 EC1-2 in all species exhibiting a higher average number of fixed mutations in non-interface sites and higher CD compared to PCDH15 EC1-2 (supplementary table S3), and with the mouse complex exhibiting a higher average number of fixed interface mutations and lower interface CD compared to the other three complexes (supplementary table S4).

## DISCUSSION

Previous studies have investigated the origin and evolution of the cadherin superfamily (Abedin and King 2008; Dickinson et al. 2011; Hulpiau and van Roy 2011; Oda and Takeichi 2011; Gul et al. 2017; Green et al. 2020). Gul et al. explored the evolutionary relationships of the various cadherin sub-types, and determined that cadherin-mediated adhesion likely existed in the last common ancestor of all animals ∼600 million years ago (Gul et al. 2017). From the generally longer ancestral cadherins, genome duplications and subsequent loss and reshuffling of EC repeats eventually led to the cadherin repertoire seen in animals today (Hulpiau and van Roy 2011; Oda and Takeichi 2011). However, no studies to date have investigated the evolution and diversification of CDH23 and PCDH15 from a biophysical and structural perspective. The data presented here, obtained using a variety of experimental and computational techniques, begin to provide a more thorough and quantitative picture of how this protein complex evolved from the ancestral state to modern vertebrates.

Our simulations and experiments reveal species-specific differences in the force response and affinity of the tip-link complex. In both all-atom and CG SMD simulations, the lizard complex exhibited the highest force required to unbind the complex compared to mouse at 0.1 nm/ns (fig. 3E, fig. 4A), a higher BSA and energy of unbinding in CG simulations (fig. 4H and J), and a higher affinity (lower *K_d_*) in SPR experiments (fig. 2). The molecular basis for this more robust complex is the result of a loss of interactions in the mammalian lineage that are formed at or near the interface of the AncAm and reptilian complexes. These interactions increase allosteric communication between CDH23 and PCDH15 in the lizard relative to the mouse (supplementary fig. S4A and D), form favorable contacts at the force peak that are less prominent in the mouse (supplementary fig. S6A-C), result in an increase in BSA relative to the mouse (supplementary fig S6D-F), and allow conserved residues to participate in intermolecular interactions only in the AncAm and lizard by freeing them from intramolecular interactions (supplementary fig. S9B). Additionally, the closer association between CDH23 EC1 and PCDH15 seen in the AncAm and lizard complex permits the formation of interactions between residues that, while present in mammals and reptiles, only form in the ancestral or reptilian complexes (supplementary fig. S9B). Similar results can be seen for the pigeon and AncAm, indicating that this is a more ancestral state of the CDH23-PCDH15 complex relative to the mammalian complex.

From an organismal point of view, the apparently tighter CDH23-PCDH15 EC1-2 interaction in the lizard did not evolve to accommodate higher frequency or higher amplitude auditory input. Lizards generally perceive sound in a frequency range of about 0.1 to 6 kHz, while mammalian hearing generally exceeds this threshold by more than an order of magnitude (Manley et al. 2004; Manley 2011). Three explanations could account for this variation: (1) selection acted on the tip-link complex of the anole lizard to accommodate some other lifestyle requirement, (2) fixation of neutral mutations resulted in this increase in unbinding forces and energy, or (3) selection acted on a different property of the CDH23-PCDH15 tip link, such as binding kinetics, that incidentally affected unbinding forces. The first explanation is unlikely, as our evolutionary analyses of the CDH23 and PCDH15 sequences detected minimal positive selection on EC1-2 of both proteins, which also casts doubt on the third explanation, though the *k_on_* values were higher in the lizard complex compared to the mouse and AncAm in SPR experiments (fig. 2B). The second hypothesis is plausible and can explain the variety of force peaks exhibited by the complexes in all vertebrate lineages (fig. 4A). Even in the absence of any external stimuli, the tip-link is under a resting tension that has been measured experimentally to range from ∼5 to 34 pN, depending on the measurement location in the cochlea (Jaramillo and Hudspeth 1993; Tobin et al. 2019). As mutations accumulate in the tip-link complex during evolution, the unbinding force will be altered and allowed to evolve stochastically. However, if one or a series of deleterious mutations reduces the unbinding force below the resting tension (or some threshold near the resting tension), it will be quickly purified from the population. Under this “minimum threshold” model of tip-link bond evolution, the neutral fixation of a series of mutations (or retention of ancestral features) in the lizard, after the mammal and reptile lineages split from the ancestral amniote, drove the unbinding force far above the lower threshold required for the anole complex. Through neutral and purifying selection, the mouse complex remained closer to the lower threshold, while still fulfilling the requirement to transduce force in the presence of an auditory stimulus.

This model can be extended to include a “maximum threshold” to the mammalian complex, which provides a physiological explanation for the lower mechanical strength exhibited by the mouse complex. The lower force of unbinding in mammals may act as a safety mechanism, in which the tip-link is selected to unbind at high frequency, high amplitude sound before the rest of the protein or hair cell is damaged. Because the surrounding inner-ear organs and tissue have not evolved to respond to high frequency sound in reptiles, this potentially damaging stimulus would not be transferred to reptilian tip links, and the tip-link complex could neutrally evolve to withstand higher forces without penalty. In this way, purifying selection could have acted to reduce and maintain the mammalian complex within a certain force range from the ancestral state, while the reptilian complexes are free to diverge neutrally, explaining the functional divergence in these proteins without the need to appeal to positive selection. This hypothesis also does not exclude the possibility of positive selection acting on these proteins at previous stages of vertebrate evolution. In fact the Selectome server (Proux et al. 2009) does predict lineage-wise positive selection to have acted in both the CDH23 and PCDH15 genes after divergence from the ancestral euteleost, but before the emergence of most extant vertebrate species. This suggests that while positive selection likely played a role in the initial evolution of these proteins in vertebrate evolution, they have been evolving primarily neutrally (or near-neutrally) since the divergence of most modern vertebrates. Additionally, while there is evidence of positive selection acting on CDH23 and PCDH25 (Shen et al. 2012; Moreira et al. 2021; Trigila et al. 2021), the sites identified are not found at the CDH23-PCDH15 interface.

This threshold model of tip-link bond evolution provides a possible explanation for the similar response to force in the lizard and mouse complex only at lower pulling speeds, where at 0.001 nm/ns the forces required to break the complex were not significantly different between the lizard and mouse (fig. 4C). At 0.001 nm/ns, the mouse complex is operating within the threshold limits of purifying selection and better reflects its physiological function. Higher speeds (0.1 and 0.01 nm/ns) are beyond this limit, and because in this model the ability to withstand these forces would be purified in the mouse as a safety mechanism, it exhibits a reduced ability to remain bound at higher forces. The lizard, on the other hand, would not experience any limit on its ability to withstand force at these pulling speeds, and as a result would maintain adhesion at higher force.

An alternative explanation for the differing response to force in these vertebrate species that is beyond the scope of this work may lie in regions of the heterotetrameric tip link beyond EC1-2. While the CDH23-PCDH15 EC1-2 dimer is ultimately responsible for force transduction, it does so in the context of other structural features that may exhibit functionally significant species-specific changes. The PCDH15 X-dimer, formed at an EC2-3 interface, and an interaction facilitated by MAD12, are two points of dimerization in PCDH15 that have functional implications upon the application of force (De-la-Torre et al. 2018; Dionne et al. 2018; Ge et al. 2018; Choudhary et al. 2020; Mulhall et al. 2021), and may have evolved to respond to different frequencies.

In rough agreement with the smooth transition from basal vertebrates to more recently evolved lineages, as is seen with maximum hearing frequency threshold and auditory papillae length, the force required to unbind the CDH23-PCDH15 EC1-2 complex shows some degree of evolutionary pattern in the results of the CG simulations (fig. 4A). For example, the turtle is believed to represent the primitive state of the tetrapod inner ear (Coffin et al. 2004), having undergone the least amount of anatomical innovations, and also exhibited the lowest average force peak and energy of unbinding of all tetrapods. Similarly, the fish inner ear is a relatively simple structure compared to land animals, as it has not been subjected to the selection pressures of air-based audition. As with the turtle, the average force peaks for the zebrafish and the rice fish were lower than most other animals (fig. 4A). However, the yellowtail complex exhibited a significantly higher force peak than the other two fish and the mouse, which could suggest an evolutionary innovation that is required for this animal’s lifestyle, or the result of neutral evolution. Finally, there is some suggestion that certain functional requirements can have an evolutionary effect on tip-link strength with the results of the whale, which exhibited the highest force peaks of the mammals (fig. 4A). Whales utilize ultrasonic echolocation to communicate, and have developed several specializations in their inner ear that allows them to hear underwater at frequencies well beyond that of most land-mammals (Ketten 1997). A stronger tip-link may be another innovation to allow this specialized form of communication in various species, including some frogs (Shen et al. 2008).

While the differences observed in the unbinding forces obtained from SMD and experimental affinities between species are small, some perturbations in the tip-link bond interface that impair but do not abolish the interaction can have significant physiological effects. For example, the PCDH15 EC1-2 fragment carrying the deafness mutation R113G still binds to CDH23 with reduced affinity in isothermal titration calorimetry experiments (Sotomayor et al. 2012), but the mechanical strength of the heterotetrameric complex is greatly reduced (Mulhall et al. 2021) indicating that drastic functional changes can result from relatively subtle molecular alterations in the tip-link complex.

In agreement with the significant differences seen in forces of unbinding in all-atom and CG-SMD simulations, and of energies of unbinding from the umbrella sampling simulations, the lizard and AncAm did exhibit slight but significant reduction in *K_d_* values in SPR experiments compared to the mouse (fig. 2). However, the differences in experimental affinities were not as drastic as the differences seen in force peaks and energies of unbinding. While a correlation between off-rates and unbinding forces has been established for some protein-ligand systems (Schwesinger et al. 2000), the relationship between equilibrium properties (*K_d_*) and force of unbinding is not necessarily clear for protein-protein interactions. In the pulling simulations with the largest force of unbinding, certain interface interactions exhibited a sudden decrease in distance and increase in stability just before the force peak (supplementary fig. S6C), while in equilibrium, these interactions were less stable (supplementary fig. S3). This effect was especially prominent in the mouse complex and demonstrates how the application of force can modulate interactions at the interface in a manner that is not observed in equilibrium. As discussed above, the tip-link is under tension and the force-dependent difference between complexes is likely to be more relevant from an evolutionary perspective, as selection on the interface will be acting in the presence of force.

The differences observed and predicted for the tip-link complexes of varying species are the result of unique evolutionary processes, and the molecular evolution simulations presented here begin to elucidate the evolutionary constraints responsible for these differences. During these simulations, the non-mammalian interfaces exhibited fewer substitutions and higher interface CD during the simulation relative to the mouse (supplementary fig. S10I-L, supplementary table S2). This result suggests that a more crowded interface is less likely to accommodate new mutations during evolution and is likely under greater evolutionary constraint, an effect that has been previously reported in different systems (Eames and Kortemme 2007; Franzosa and Xia 2009). Because of this, less dense interfaces or residues at the periphery of interfaces may tolerate mutations more and may even be more likely to facilitate evolutionary innovations. Evidence for this is seen in the constant force simulations in which a substitution in the mouse prevents an interaction in the interface that is present in AncAm, lizard, and pigeon (supplementary fig. S9B). The decreased interface CD seen in the mouse in both MD and evolutionary simulations suggests that it may have decreased during evolution from the ancestral state to the mouse, permitting substitutions at or near the interface that result in the lower affinity and force response observed for the mouse complex in MD simulations and SPR.

Beyond the interface, results of the evolutionary simulations demonstrate that the three-dimensional structure of CDH23 and PCDH15 can constrain their rate of evolution and influence the functional properties of the extant homologs. In all species, CDH23 accumulated a greater number of fixed mutations per residue compared with PCDH15 during the simulations (supplementary fig. S10E-H, supplementary fig. S11, supplementary table S1), and maintained higher CD throughout the evolutionary trajectories (supplementary fig. S10A-D). This provides a possible biophysical basis for the higher average overall evolutionary rate of total sites, higher average rate of buried sites, and lower sequence identity observed for CDH23 compared to PCDH15 and suggests that PCDH15 EC1-2 is under greater evolutionary constraint compared to CDH23 EC1-2. This conclusion is further supported by the BUSTED analysis. This higher rate of evolution in CDH23 EC1-2 may be partly responsible for observed changes in affinity and interaction strength, as all three substitutions that modified the response to force in the mouse complex in all-atom MD (fig. 3F-I) were found in solvent exposed sites of CDH23, and all are predicted by FoldX to destabilize the protein fold if inserted in the AncAm CDH23 EC1-2 structure. The increased CD of CDH23 EC1-2 may facilitate such destabilizing but function altering mutations. Evidence for this phenomenon has been previously reported for other systems (Faure and Koonin 2015). These results demonstrate how fundamental biophysical properties of the tip link and its interaction could have shaped its evolutionary trajectory, the result of which gave rise to the divergent tip-link function observed in diverse extant species.

The observed correlation between the differing biophysical properties and rate of evolution between CDH23 and PCDH15 EC1-2 suggests an intriguing fundamental explanation for the increased presence of deafness-causing mutations found in CDH23 relative to PCDH15. In the evolutionary simulations, CDH23 EC1-2 exhibited a marked increase in substitutions that was irrespective of species, type of site (buried or interfacial), or the chemical properties of the residue being inserted (supplementary fig. S11). Supported by the results of the evolutionary sequence analyses, this suggests that CDH23 EC1-2 is more robust to mutations that would have a destabilizing effect on its structure or interface and would more readily accommodate fitness-reducing but non-lethal mutations that result in deafness. This effect has been previously identified, in which relaxed evolutionary selection in certain genes was responsible for the propagation of disease-causing variants in different vertebrate species (Cui et al. 2019). In the context of the tip link, the results presented here provide a possible biophysical basis for this phenomenon and the apparent preference for CDH23 of deafness-causing variants (Jaiganesh et al. 2017).

The results presented here demonstrate that the rate and pattern of the molecular evolution of CDH23 and PCDH15 are distinct from one another, constrained to some degree by their three-dimensional structure, and coupled to one another through an interaction that is responsible for force transduction in the inner ear of vertebrates. Overall CD and decreased interface CD are two biophysical properties that were revealed by evolutionary simulations to permit or constrain evolutionary innovation. The biophysical effect of these differential evolutionary forces is evident in MD simulations and in SPR experiments, which demonstrate small but significant species-level differences in the CDH23-PCDH15 interface both in equilibrium and under applied force as a result of sequence level evolutionary divergence and offers a glimpse into how the inner ear of these different animals evolved. To what extent these effects extend to other members of the cadherin superfamily or to other families of force-conveying protein interactions is unclear. Additionally, there are undoubtedly several evolutionary forces beyond those explored here that influence the evolution of these proteins, such as epistasis (Starr and Thornton 2016; Miton et al. 2021), allosteric effects (Raman et al. 2016), and the evolution of posttranslational modification sites (Zielinska et al. 2012; Landry et al. 2014; Kim and Hahn 2015). Finally, while the tip link is known to exist as a heteroteramer in vivo (Kazmierczak et al. 2007; Choudhary et al. 2020), the focus of this work was on the heterodimeric interaction, as it is this interface that is responsible for transducing force in the inner ear. How the overall heterotetrametric complex evolved to respond to force in other species remains unknown.

## MATERIALS AND METHODS

### Ancestral Reconstruction

*CDH23* and *PCDH15* genes were identified and downloaded from the UCSC genome browser 100 way MultiZ alignments (https://hgdownload.soe.ucsc.edu/goldenPath/hg38/multiz100way/). Genes from human (hg38), chimpanzee (panTro4), rhesus macaque (rheMac3), crab-eating macaque (macFas5), olive baboon (papAnu2), green monkey (chlSab2), white-tufted-ear marmoset (calJac3), Bolivian squirrel monkey (saiBol1), northern white-cheeked gibbon (nomLeu3), orangutan (ponAbe2), small-eared galago (otoGar3), Chinese tree shrew (tupChi1), pig (susScr3), Tibetan antelope (panHod1), sheep (oviAri3), goat (capHir1), cow (bosTau8), cat (felCat8), dog (can Fam3), Pacific walrus ( odoRosDiv1), Weddell seal (lepWed1), ferret (musFur1), panda (ailMel1), white rhino (cerSim1), horse (equCab2), black flying fox (pteAle1), megabat (pteVam1), prairie vole (micOch1), Chinese hamster (criGri1), golden hamster (mesAur1), mouse (mm10), rat (rn6), star-nosed mole (conCri1), alpaca (vicPac2), Bactrian camel (camFer1), hedgehog (eriEur2), Cape elephant shrew (eriEur2), aardvark (oryAfe1), manatee (triMan1), David’s myotis bat (myoDav1), big brown bat (eptFus1), microbat (myoLuc2), elephant (loxAfr3), lesser Egyptian jerboa (jacJac1), naked mole rat (hetGla2), guinea pig (cavPor3), chinchilla (chiLan1), brush tailed rat (octDeg1), Cape golden mole (chrAsi1), shrew (sorAra2), rabbit (oryCun2), pika (ochPri3), dolphin (turTru2), killer whale (orcOrc1), squirrel (speTri2), armadillo (dasNov3), tenrec (echTel2), opossum (monDom5), Tasmanian devil (sarHar1), wallaby (macEug2), Saker falcon (falChe1), Peregrine falcon (falPer1), budgerigar (melUnd1), scarlet macaw (araMac1), Puerto Rican parrot (amaVit1), rock pigeon (colLiv1), collared flycatcher (ficAlb2), zebra finch (taeGut2), Tibetan ground jay (pseHum1), medium ground finch (geoFor1), white throated sparrow (zonAlb1), chicken (galGal4), turkey (melGal1), mallard duck (anaPla1), American alligator (allMis1), green sea turtle (cheMyd1), painted turtle (chrPic2), Chinese softshell turtle (pelSin1), spiny softshell turtle (apaSpi1), lizard (anoCar2), and platypus (ornAna1) were included in analyses. Amino acid sequences were aligned using either MUSCLE or the MAFFT (Katoh et al. 2002) algorithm implemented in GUIDANCE2 (Sela et al. 2015). Alignment uncertainty was estimated with 1000 bootstrap replicates and sites with scores less than 0.8 masked from downstream analyses (source data 1 tables 1 and 2, source data 2 tables 1 and 2). Both methods generated the same alignment, therefore the masked MAFFT alignment was used for ancestral reconstruction. Ancestral CDH23 and PCDH15 proteins were inferred using the ASR module of the Datamonkey web-sever (Delport et al. 2010) using the alignment, the best fitting model of amino acid substitution (inferred to be JTT with model defined amino acid frequencies for both proteins; source data 1 table 3 and source data 2 table 3), site-to-site rate variation accommodated with a general discrete distribution with 3 rate classes, and the species phylogeny. The ASR module implements three methods to reconstruct ancestral sequences (joint, marginal, and sampled) (Kosakovsky Pond and Frost 2005), all three methods reconstructed the same ancestral sequences (source data 1 tables 4 and 5, source data 2 tables 4 and 5). IQTREE was used as an additional method for comparing reconstructions. The AncMam and AncAm sequences of CDH23 and PCDH15 were inferred by first reconstructing a tree by IQTREE and using this tree for the reconstruction. Similarly to Datamonkey, the JTT model of nucleotide substitution was used. The resulting sequences agreed with sequences inferred by Datamonkey. All alignments and ancestral reconstructions are provided in supplementary dataset 1.

### Protein Production and Purification

All constructs used for crystallography and as ligands in SPR experiments contained a hexahistidine tag and were subcloned into *NdeI* and *XhoI* sites of the pET21a vector. These included the AncMam CDH23 EC1-2 (all residues in fig. 1E), AncAm CDH23 EC1-2 (all residues in fig. 1E), lizard CDH23 EC1-2 (NCBI Ref. XP_016847668.1, residues N3-P211), turtle CDH23 EC1-2 (NCBI Ref. XP_007054748.1, residues N3-P211), AncAm PCDH15 EC1-2 (all residues in fig. 1E), mouse PCDH15 EC1-2 (NCBI Ref. XP_017169273.1, residues Q1-D233), and lizard PCDH15 EC1-2 (NCBI Ref. XP_016851436.1, residues Q1-P242). These residue ranges were chosen to start immediately after the signal peptide at the N-terminus as predicted by SignalP (http://www.cbs.dtu.dk/services/SignalP/), and to end before the start of EC3 at the C-terminus based on the conserved DXNDN Ca^2+^-binding motif following the numbering as in fig. 1E. These constructs were expressed in BL21CodonPlus(DE3)-RIPL cells and cultured in 2 L of Terrific Broth (TB). Induction was initiated with 200 µM of IPTG at OD_600_ ∼0.4 to 0.6 and cells were cultured at 37 °C for ∼16 h. Cells were then pelleted and lysed in denaturing buffer with 20 mM TrisHCl (pH 8), 6 M guanidine hydrochloride (GuHCl), 10 mM CaCl_2_ and 20 mM imidazole. The cell lysate was centrifuged at 20,000 rpm, and the supernatant was nutated with 6 mL of Ni-Sepharose beads (GE Healthcare). The protein fragment of interest was eluted with denaturing buffer supplemented with 500 mM imidazole. The PCDH15 fragments for all species were refolded using one of two methods: the first was a six-step dialysis method, in which 2 mM dithiothreitol (DTT) was added to the protein, which was then dialyzed using MWCO 2,000 membranes against buffer containing 6 M, 3 M, 2 M, 1 M, 0.5 M, and 0 M GuHCl and 400 mM arginine for 24 hours (6 to 1 M) and 12 hours (0.5 and 0 M, both of which also contained 350 µM glutathione oxidized). The second refolding method used for PCDH15 fragments employed a drop-by-drop dilution approach, in which ∼40 mL of elution was dropped at ∼10 drops/min into a large (1 L) reservoir of buffer. For all CDH23 constructs, an overnight dialysis method was employed. In all cases, for both CDH23 and PCDH15, the buffer contained 20 mM TrisHCl (pH 8.0), 150 mM KCl, 400 mM arginine and 5 mM CaCl_2_, in addition to the associated GuHCl as indicated for the stepwise buffers.

All constructs encoding for the protein fragments that were used as the tagless analyte in SPR experiments were produced using the IMPACT protein purification kit (NEB) following the manufacturer’s guidelines for cloning and refolding from denaturing conditions. These included AncAm CDH23 EC1-2, mouse CDH23 EC1-2 (NCBI Ref. XP_030100929.1, residues Q1-D205), and lizard CDH23 EC1-2, which were subcloned into the pTXB1 vector using *NdeI* and *Sap1* restriction sites, which results in a self-cleaving intein tag fused to the C-terminus of the target construct. These constructs were expressed in BL21CodonPlus(DE3)-RIPL cells and cultured in 2 L of TB. Induction was initiated with 200 µM of IPTG at OD_600_ ∼0.4 to 0.6 and expression was performed at 37 °C for ∼16 h. The purification was performed as described previously (Choudhary et al. 2017), with some modifications. Briefly, cells were lysed in denaturing conditions with 20 mM TrisHCl (pH 8), 6 M GuHCl, 10 mM CaCl_2_ and 20 mM imidazole. The lysate was centrifuged at 20,000 rpm, and the supernatant was dialyzed against buffer containing 8 M, 6 M, 4 M, 2 M, and 0 M urea for ∼24 hours each, where the final two buffers also contained 0.1 mM oxidized glutathione and 1 mM reduced glutathione. This was then loaded and incubated in a gravity column containing 25 mL of chitin bead slurry (NEB), then washed with 400 mL column buffer (20 mM TrisHCl pH 8, 500 mM NaCl, 2 mM CaCl_2_), followed by incubation with cleavage buffer (60 mL column buffer with 50 mM DTT) for 48 h. This was then eluted with 40 mL of column buffer. All proteins were purified using a Supderdex200 or Superdex75 size exclusion chromatography column in 20 mM TrisHCl (pH 8.0), 150 mM KCl and 2 mM CaCl_2_. PCDH15 108N mutations were generated using the QuikChange Lightning mutagenesis kit (Agilent) and mutant protein fragments were expressed and purified identically to the wild-type constructs. All DNA constructs were sequence verified.

### Crystallization and Structure Determination

The sitting-drop vapor diffusion method was used to grow crystals from a 1:1 mix of protein and reservoir solution at 4 °C. Buffer conditions and cryoprotection buffers were prepared as indicated in supplementary table S5. Crystals were cryo-cooled in liquid N_2_, and X-ray diffraction data sets were collected as indicated in supplementary table S6. All datasets were processed with HKL2000 (Otwinowski and Minor 1997) and structures determined by molecular replacement using PHASER (McCoy et al. 2007). For all constructs, the crystal structure of the mouse CDH23 EC1-2 (PDB ID 2WHV) (Sotomayor et al. 2010) was used as a template for molecular replacement. COOT (Emsley and Cowtan 2004) was used for model building, and restrained refinement was performed with REFMAC5 (Murshudov et al. 2011). Data collection and refinement statistics for all structures are provided in supplementary table S6. Before deposition, all structures were analyzed using Procheck (Laskowski et al. 1993), Whatcheck (Hooft et al. 1996), and Checkmymetal (Zheng et al. 2014).

### SPR Experiments

SPR experiments were performed using an OpenSPR instrument and NTA sensor chips from Nicoya Lifesciences. For all species, CDH23 was chosen as the tagless analyte, with PCDH15 as the bound ligand. Functionalization of PCDH15 to the NTA chip was achieved via the 6X-His tag of PCDH15 and following the manufacturer’s guidelines. Injection of CDH23 was performed at 35 µl/min in a series of 10 concentrations: 25, 20, 18, 15, 12, 10, 8, 5, 2 and 1 µM. A sample loop of 100 µl was used with a flow rate of 35 µl/min. The signal was given sufficient time to reach baseline before subsequent injection, typically 200-400 seconds. All samples used identical buffers from size exclusion chromatography experiments. Data were reference and buffer subtracted and subsequently analyzed using the TraceDrawer software (Nicoya) using a kinetic evaluation, assuming a 1:1 binding interaction. In rare instances, the signal for an injection appeared to be higher or lower than expected as compared to previous or subsequent injections. In such cases, these injections were excluded from data analysis. If data collection was interrupted for a certain injection, for example due to the presence of air bubbles in the flow cell, that concentration was excluded from analysis. All reported *K_d_* values were calculated with curves from at least 7 concentrations included in the theoretical fit to the data. Fits were only accepted when errors of the resulting *k_on_*, *k_off_*, and *K_d_* values were at least two orders of magnitude smaller than the reported values.

### All-Atom MD Simulations

The structure of the mouse CDH23-PCDH15 complex (PDB ID 4APX) (Sotomayor et al. 2012) was used both to simulate the mouse complex, and as the base for homology models of the pigeon, AncAm, and lizard all-atom complexes. In the case of these non-mammalian complexes, *in silico* amino acid substitution was performed in COOT (Emsley et al. 2010) to change the mouse sequence (NCBI Ref. XP_030100929.1, residues Q1-Q207 for CDH23, NCBI Ref. XP_017169273.1, residues Q1-L235 for PCDH15) in 4APX to pigeon (NCBI Ref. XP_021148560.1, residues N3-D205 for CDH23, and NCBI Ref. XP_021147002.1, residues H1-L235 for PCDH15), AncAm (fig. 1E), and lizard (NCBI Ref. XP_016847668.1, residues N3-Q207 for CDH23, and NCBI Ref. XP_016851436.1, residues Q1-L235 for PCDH15). All sequences were cut at the same location in the alignment with the exception of CDH23 in the mouse, which was predicted to contain two extra residues at its N-terminus, and CDH23 in the pigeon, which was simulated with two fewer residues at its C-terminus. Clashes were removed and arranged in a biochemically reasonable manner after substitutions were introduced using the real-space refinement feature of COOT. Because the CDH23 EC1-2 structures of such diverse vertebrates as fish, lizard, and mouse show a remarkable degree of structural conservation despite their sequence divergence (fig. 1F, supplementary figure S2A, D, G and J), these serve as reasonable homology models for further simulation. The VMD psfgen, solvate, and autoionize plugins were used to build all systems (Humphrey et al. 1996). Water boxes were made large enough to accommodate unbound states, and 150 mM KCl was added to all systems to imitate the physiologically high concentration of potassium in the inner ear endolymph (Bosher and Warren 1968). The N-termini of all proteins were capped with -NH^3+^ (NH3 CHARMM atom type), and the C-termini with a -COO^-^ carboxylate group (CC and OC CHARMM atom types).

Simulations were ran using NAMD 2.12 (Phillips et al. 2005) with the CHARMM36 force field, and the TIP3P explicit model of water (Huang and MacKerell 2014). A cutoff of 12 Å was used for van der Waals interactions, and long range electrostatic forces were computed using the Particle Mesh Ewald method with a grid point density of >1 Å^−3^. The SHAKE algorithm was used with a timestep of 2 fs. The *NpT* ensemble was used at 1 atm with a hybrid Nosé-Hoover Langevin piston method, a 200 fs decay period, and with a 100 fs damping time constant. All systems were minimized for 5000 steps, followed by 100 ns of equilibration. SMD simulations were initiated from different starting configurations throughout the equilibrium simulations (see Results). Constant velocity stretching was applied using the SMD method (Izrailev et al. 1998; Grubmüller 2005; Lee et al. 2009; Rico et al. 2013; Valotteau et al. 2019; Franz et al. 2020) and through the NAMD Tcl interface. A virtual spring with a stiffness of *k_s_* = 1 kcal mol^−1^ Å^−2^ was attached to the C-terminal C_α_ carbon atoms of each chain, and each was then stretched at a constant velocity of 0.05 nm/ns. Forces were calculated based on the extension of the spring. For the constant force pullings, a constant force of 150 pN was applied to the C-terminal C_α_ carbon of PCDH15 using SMD forces in NAMD, with the C-terminal C_α_ carbon of CDH23 constrained (*k_s_* = 1 kcal mol^−1^ Å^−2^). Equations were fit to simulation data after end-to-end distance was normalized by setting the initial distance before force was applied to zero.

For SMD simulations, end-to-end distance was measured from the position of the C-terminal C_α_ atom of one monomer and the C-terminal C_α_ atom of the other. Spring constant and damping coefficient were estimated by fitting a model of a simple spring in a viscous environment to the constant-force simulation data. Force peaks for all-atom simulations were taken from 50 ps running averages of the force plots to eliminate local fluctuations. CD was measured as the average number of inter-residue contacts per residue in the selection of residues being measured. While the cutoff value used to measure inter-residue contacts varies in the literature, here a contact was recorded if each residue’s C_α_ atom were within 9 Å of one another (Liao et al. 2005; Yeh et al. 2014), excluding contacts from residues immediately adjacent in the primary structure. Interface CD was calculated in an identical manner, but inter-protein contacts were only counted from residues that were within 6 Å of any non-hydrogen atom of a residue in the associated chain’s binding partner. For all simulations, residues were considered buried by first identifying all residues that made more than 10 contacts with other residues based on the distance of their C_α_ carbons, with further inspection of these residues to determine if their sidechains were facing the core. Any residue whose sidechain was facing the solvent was not considered buried. Residues were considered in the interface if any atom was within 6 Å of any atom of any residue in the associated chain’s binding partner, chosen to include residues that were directly involved in the complex and those on the periphery that may influence interface dynamics and association, and based on previous work (Eames and Kortemme 2007). Because the Ca^2+^-binding linker region between EC repeats was excluded from evolutionary simulation analyses due to complete conservation of these residues in sequence alignments, contact density of CDH23 EC1-2 and PCDH15 EC1-2 was calculated by averaging the contact density of EC1 and EC2 separately, excluding the linker residues. Values for contact density during the MD simulations were taken from averages of the normal distribution fit to the distribution of running averages over the course of the trajectories. Buried surface area was computed in VMD by measuring solvent accessible surface area (SASA) for CDH23 and PCDH15 separately and by subtracting SASA for the complex and multiplying by 0.5, with BSA for interacting molecules A and B computed as ½(SASA(A) + SASA(B) – SASA(AB)). Xmgrace was used to generate plots and the molecular graphics program VMD (Humphrey et al. 1996) was used to analyze trajectories.

### CG MD Simulations

As with the all-atom systems, the CG systems were created by first performing amino-acid substitution in COOT on the crystal structure of the mouse CDH23-PCDH15 EC1-2 complex (PDB ID 4APX) (Sotomayor et al. 2012) to change the sequence to the whale (NCBI Ref. XP_023987875.1 residues Q1-D205 for CDH23, XP_023977067.1 residues Q1-L235 for PCDH15), bat (XP_024416963.1 residues Q1-Q207 for CDH23, XP_024416857.1 residues Q1-L235 for PCDH15), bearded dragon (XP_020638767.1 residues N3-D207 for CDH23, XP_020662251.1 residues Q1-V234 for PCDH15), turtle (XP_014428215.1 residues N3-Q207 for CDH23, XP_014427543.1 residues Q1-L235 for PCDH15), alligator (XP_019339897.1 residues N3-Q207 for CDH23, XP_019339843.1 residues Q1-L235 for PCDH15), and three different bony fish, including *Danio rerio* (zebrafish) (NP_999974.1 residues N3-Q207 for CDH23, XP_009305800.1 residues Q1-V234 for PCDH15), *Oryzias latipes* (rice fish) (XP_023819152.1 residues S2-Q207 for CDH23, XP_023818757.1 residues Q1-V234 for PCDH15), and *Seriola lalandi dorsalis* (yellowtail) (XP_023267596.1 residues S2-Q207, XP_023282391.1 residues Q1-V234 for PCDH15). The AncAm, anole lizard, and pigeon had already been created for the all-atom simulations and are described above. Real space refinement in COOT was used to remove clashes and rearrange the substituted residues in a biochemically reasonable configuration. The resultant all-atom structures were converted to coarse-grain models using the martinize.py script available on the martini webpage (http://cgmartini.nl/index.php/tools2/proteins-and-bilayers/204-martinize). An elastic network model, in which harmonic bonds are added between beads to ensure the secondary structure is maintained during simulation, was implemented through the “-elastic” flag of the martinize.py script. Force constants of 500 kJ mol^-1^ nm^-2^ were chosen for the elastic network based on comparison to all-atom simulations of cadherins. Ca^2+^ ions were represented as Qd type beads with a +2 charge. Each system was solvated in a waterbox with enough room for forced unbinding using the Gromacs solvate command. Water was represented as P4 bead types, and the system was ionized with 150 mM NaCl represented as Qd bead types with a +1 charge and Qa bead types with a −1 charge, respectively.

All CG simulations were ran using the Gromacs MD engine (Abraham et al. 2015) and the Martini force field (Monticelli et al. 2008) with a 9 fs timestep. Each system was minimized for 50,000 steps, followed by backbone-constrained equilibration in which constraints were reduced sequentially from 4,000, to 2,000, 1,000, and 500 kJ mol^-1^ nm^-2^ for 100,000 steps each. Finally, a free equilibration was ran for 2,225,000 steps (∼20 ns) with no constraints. Constant-velocity pulling was implemented using the Gromacs “pull” code, in which a harmonic potential with a stiffness of 418.4 kJ mol^-1^ nm^-2^ was applied to the C-terminal backbone beads of each chain at 0.05 nm/ns, unless otherwise indicated. Force peaks for CG simulations were taken from 1 ns running averages, chosen because of the higher degree of noise inherent in CG data. BSA for CG simulations was computed as described for all-atom simulations.

To calculate the potential of mean force (Park and Schulten 2004) for selected species, initial conformations were saved every 3 ns during the first SMD simulation for each system. If sufficient overlap of umbrella sampling windows was not achieved, additional starting configurations were saved. From these starting configurations, constrained simulations were ran in which the C-terminal backbone bead of each chain was held fixed for 11,150,000 steps (∼100 ns). From these sampling windows, the WHAM method as implemented in Gromacs (Hub et al. 2010) was used to construct a potential of mean force, and thus the *ΔG*, for the unbinding pathway of each complex.

### Molecular Evolution and Origin-Fixation Simulations

All simulation and analysis scripts were written in python, based on the methods of the evolutionary model outlined previously (Kachroo et al. 2015; Teufel and Wilke 2017). For each system (lizard, AncAm, mouse, and pigeon), conformations of CDH23 EC1-2, PCDH15 EC1-2, and the complex were saved as separate coordinate files taken from the starting point of the all atom MD simulations. Each was minimized using the RepairPDB command of the FoldX suite (Schymkowitz et al. 2005), and the resulting structures were used as the starting structure (wild type) in subsequent simulations. Starting from these initial structures, at each timestep, CDH23 or PCDH15 (chosen randomly) are mutated at a single site to a different amino acid, also chosen at random. Mutations are inserted using the FoldX BuildModel command, from which the energetic effect of the mutation on protein stability (*ΔΔG*_s_) is obtained. The same mutation is then introduced to the complex using the FoldX AnalyseComplex command to determine the energetic effect of the mutation on the interaction, and from which a binding energy is obtained (*ΔG*_I_).

Energy values for stability and binding energy (*ΔΔG*_s_ and *ΔG*_I_) were used to determine the fitness of the mutation based on the fitness function from Kachroo et al. 2015 with a β value of 0.1:

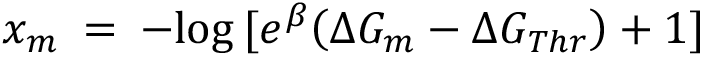

Where *x_m_* is the fitness value of the mutant, *ΔG_m_* is the energy value of the mutant (either *ΔΔG*_s_ or *ΔG*_I_), and ΔG_Thr_ is the threshold value that determines the threshold of the sigmoidal curve. The threshold was chosen as half the value of the *ΔG* (for calculating the fitness effect on stability) or half the value of binding energy (for calculating the fitness effect on binding energy) of the initial, starting structure. As in Kachroo et al. 2015, fitness values for stability and interaction were summed to give an overall fitness value for the current mutation.

To simulate an evolutionary trajectory, each step of the simulation corresponds to the insertion of a single mutation. Before proceeding to the next step, each mutation is accepted or rejected according to the Metropolis criterion with a probability distribution described previously with an effective population size *N* = 10 (Kohn et al. 2005; Kachroo et al. 2015; Teufel and Wilke 2017). If the mutation is accepted, this structure is saved, and will be the starting structure for the next mutation. If it is rejected, the process is repeated from the previous starting structure. This process was repeated until 3000 mutations had been fixed for a single simulation, corresponding to an extended evolutionary trajectory to observe the evolutionary steady-state behavior of the complex, or until 75 mutations had been fixed, corresponding roughly to the age of the vertebrate lineage. For each system, 10 independent simulations were completed, with 10 additional simulations for each system ran for 75 steps. Analysis of the simulations ran for 3000 steps was performed on the initial 10 simulations, while analysis for the shorter-timescale simulations was performed on the first 75 steps of the first 10 simulations plus the 10 simulations ran for only 75 steps for a total of 10 long-timescale and 20 short-timescale simulations for each system. Structure coordinates of fixed mutants were saved at each step for analysis of structural characteristics. Analyses of these structural features throughout the trajectory were calculated by averaging the value at each step of the simulation over all 10 simulations for the long-timescale simulations, and over all 20 simulations for the shorter-timescale simulations. Determination of CD, interface CD, and buried vs. interface residues for the evolutionary simulations was performed in an identical fashion to the all-atom MD simulations. Non-interface residues were all residues excluding interface residues, as determined in an identical manner to the all-atom MD simulations. Values for CD during the evolutionary trajectories were taken from averages of the normal distribution fit to the distribution of running averages over the course of the trajectories. Simulations that were analyzed by weighting each accepted mutation by number of attempted mutations were performed in an identical manner in 20 independent runs for each complex and were ran to 312 attempted mutations.

### Evolutionary Rate Analysis

To further characterize the molecular evolution of CDH23 and PCDH15, the site-wise rate of evolution was analyzed using the protocols outlined in Wilke et al. 2017, using identical parameters, methods, and software. Sequences were curated by hand from a variety of vertebrate species ranging from sharks and bony fish to representatives of all major groups of tetrapods, and only sequences with a complete EC1-2 of both CDH23 and PCDH15 were selected. A total of 50 sequences were used for CDH23 and 55 for PCDH15. Multiple sequence alignments were generated with MAFFT (Katoh et al. 2002), and subsequent phylogenetic inference and tree generation was performed with RAxML (Stamatakis 2014). HyPhy’s likelihood estimation of individual site rates (LEISR) method (Spielman and Pond 2018) was used to infer site-wise rates at an amino-acid level. The same sequence alignments were used to calculate sequence identity and similarity (Garcia-Boronat et al. 2008). Finally, HyPhy’s BUSTED method (Murrell et al. 2015), along with the BUSTED-compare.bf batch script (https://github.com/veg/hyphy-analyses/tree/master/BUSTED-compare), was used to test the difference in rate distributions between CDH23 and PCDH15 EC1-2 using a LRT.

## Statistical Analyses

Statistical significance for SPR results were only reported if there was a significant difference in *K_d_* values determined from estimation statistics with a 95% bootstrap confidence interval (Ho et al. 2019), which allows statistical comparison of the means of two groups with different *N*. For all other analyses in which the *N* is the same between groups, including those for MD simulations and evolutionary simulations, statistical significance was only reported if two populations exhibited a significant difference both when evaluated using a pair sampled t-test for means with a hypothesized difference of zero and a two-tailed *P* value of < 0.05, and a significant difference determined from estimation statistics with a 95% bootstrap confidence interval.

## Author Contributions

C.R.N. prepared, ran, and analyzed MD and evolutionary simulations, and wrote the manuscript. C.R.N. and D.C. performed and analyzed SPR experiments. C.R.N. and Y.N. performed crystallography experiments. V.J.L. and J.D.B. performed ASR. M.S. supervised training and edited the manuscript. V.J.L., M.S., Y.N. and C.R.N. conceived the project.

## Acknowledgements

This work was supported by the Ohio State University and by the Human Frontier Science Program (RGP0056/2018). Simulations were performed using the PSC-Bridges (XSEDE MCB140226) and OSC-Owens supercomputers (PAS1037/PAA0217). C.R.N. was supported by an OSU/NIH molecular biophysics training grant (TG32GM118291). M.S. was an Alfred P. Sloan fellow (FR-2015-6794). We thank Lahiru Wimalasena for developing the CG MD protocol used, Hong-Duc Phan for initial cloning of lizard constructs, and Colin Klaus for insightful discussions.

## SUPPORTING INFORMATION

**FIGURE S1.**
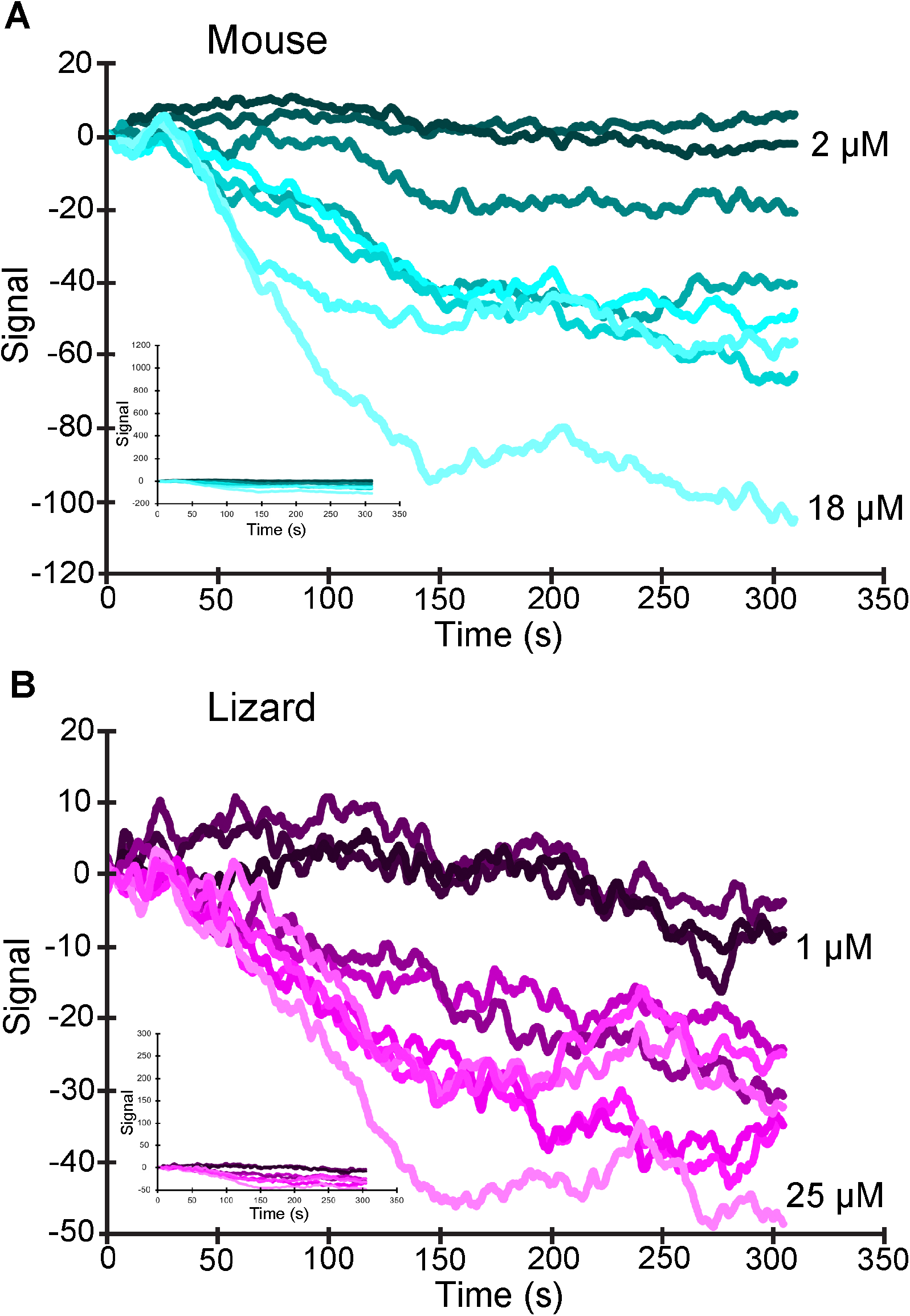
The I108N mutation in PCDH15 abolishes the tip link interaction in SPR. Raw data curves from SPR experiments performed on the mouse (*A*) and lizard (*B*) complex, in which the PCDH15 I108N mutation was present for both species. For both species, no interaction with CDH23 was observed. Insets show data using the scales used in Fig. 2.

**FIGURE S2.**
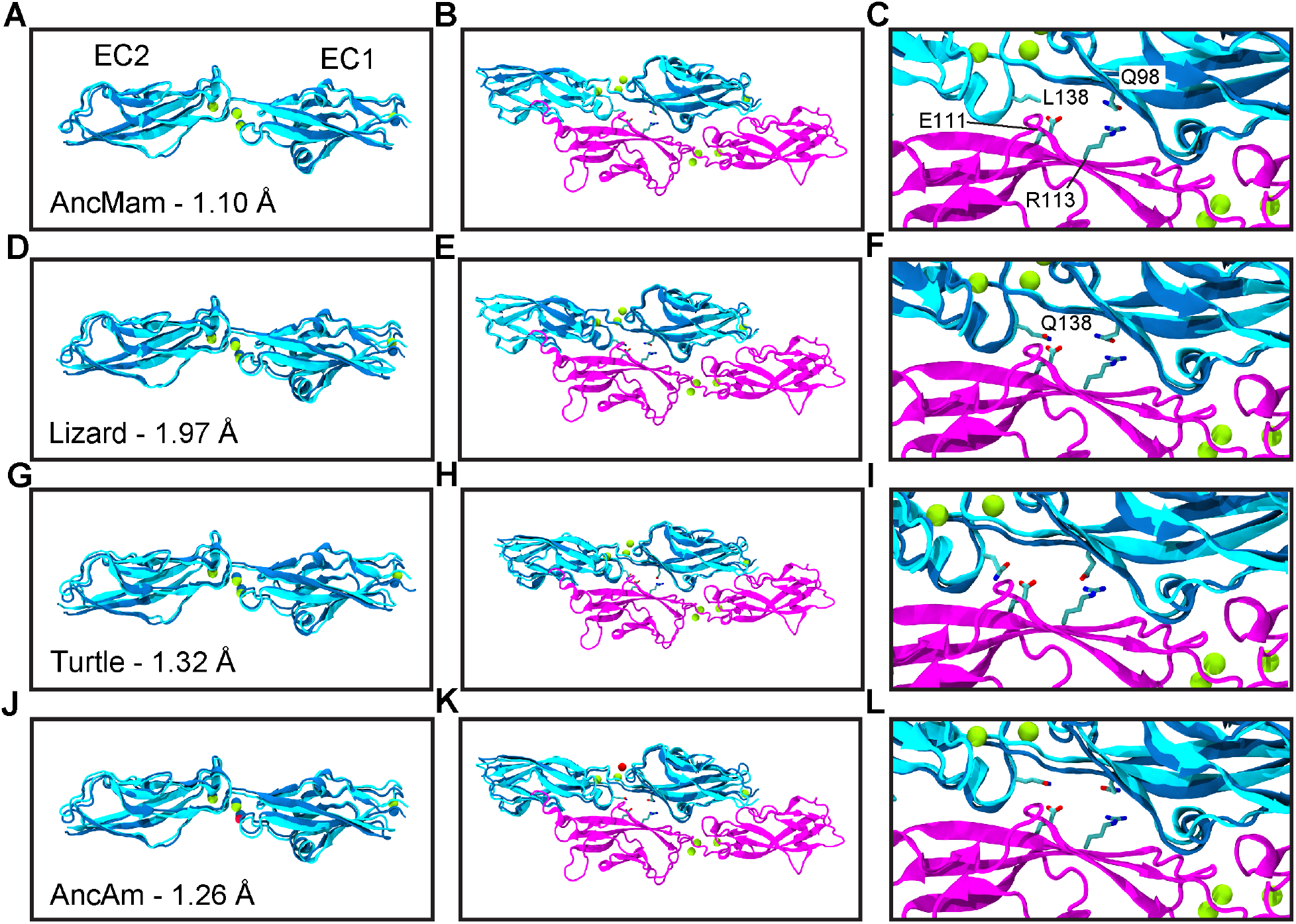
Crystal structures of CDH23 EC1-2 reveal high structural similarity in non-mammals. (*A, D, G,* and *J*) The crystal structure of the respective animal is shown in cyan aligned to the mouse CDH23 EC1-2 (PDB ID 2WHV) shown in blue. RMSD values between the structures are indicated, calculated from alignment to each structure’s C_α_ atoms. Ca^2+^ ions from the mouse are in blue, while Ca^2+^ ions from the aligned structure are in yellow. Na^+^ is shown in red. (*B, E, H*, and *K*) The crystal structure of the respective animal is shown in cyan, aligned to the mouse CDH23 EC1-2 in the crystal structure of the tip-link complex (PDB ID 4APX). The aligned structure is shown in cyan, and the mouse CDH23 EC1-2 is shown in blue. (*C, F, I*, and *L*) A closer view of the aligned complexes is shown, with specific interactions shown in licorice representation. For all complexes, residues from CDH23 are shown from the aligned structure, while residues from PCDH15 are from the original mouse structure. By alignment of the crystal structures, these important interactions are formed in the interface for all species, except for the AncMam complex, which experienced a CDH23 Q138L substitution during evolution.

**FIGURE S3.**
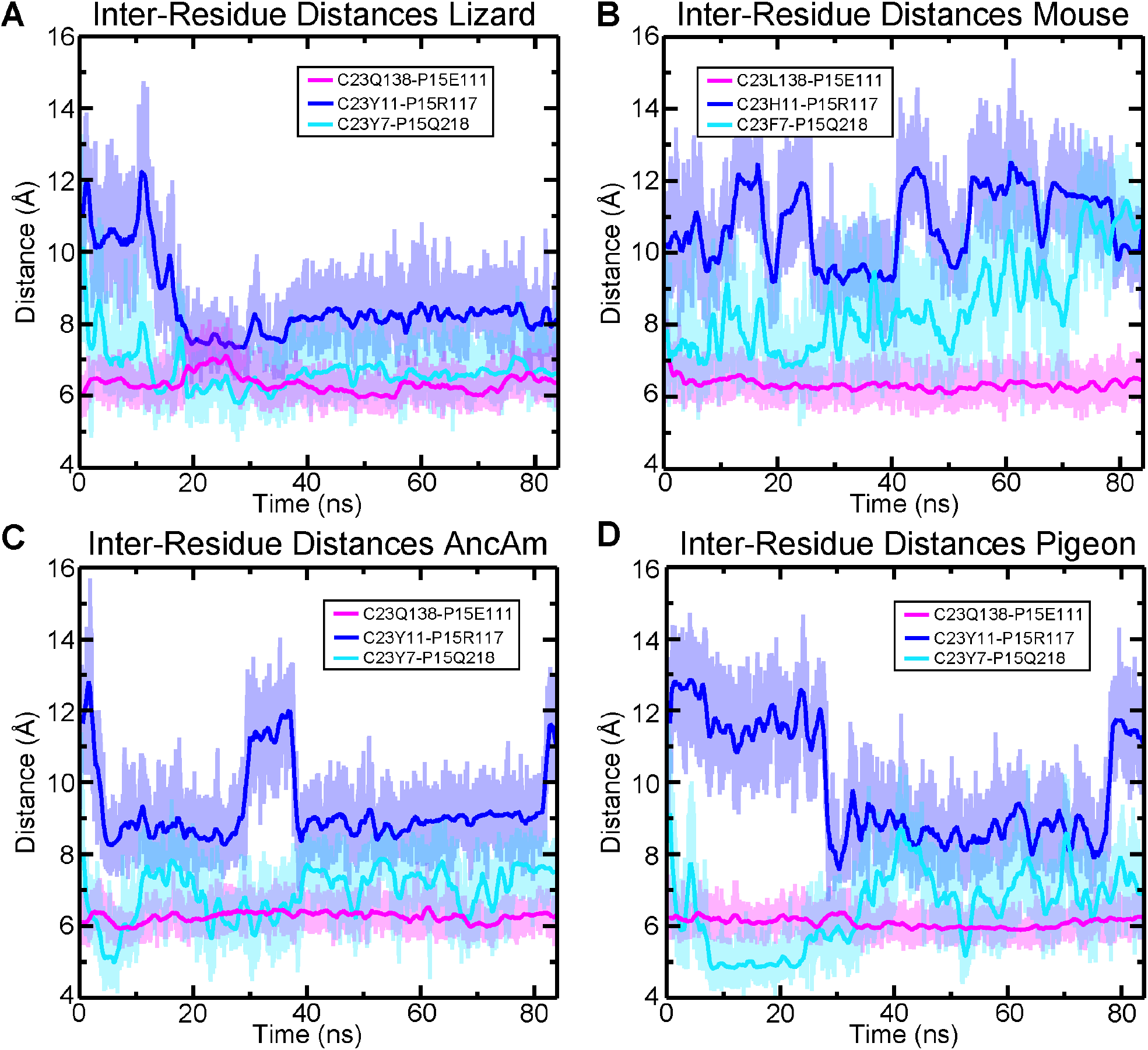
Interface dynamics in equilibrium differ between species due to evolutionary sequence changes. (*A*) Lizard interactions C23Q138-P15E111, C23Y11-P15R117, and C23Y7-P15Q218 are the most stable during all-atom MD equilibrations (Sim1-4a). (*B*) The mouse experienced substitutions during evolution that prevent these interactions from forming stably. (*C*) These interactions were present in the ancestral amniote, suggesting the mammalian lineage represents a more derived state of the complex, while the reptilian complexes are more ancestral. (*D*) As with the lizard, the pigeon complex exhibited stable interactions.

**FIGURE S4.**
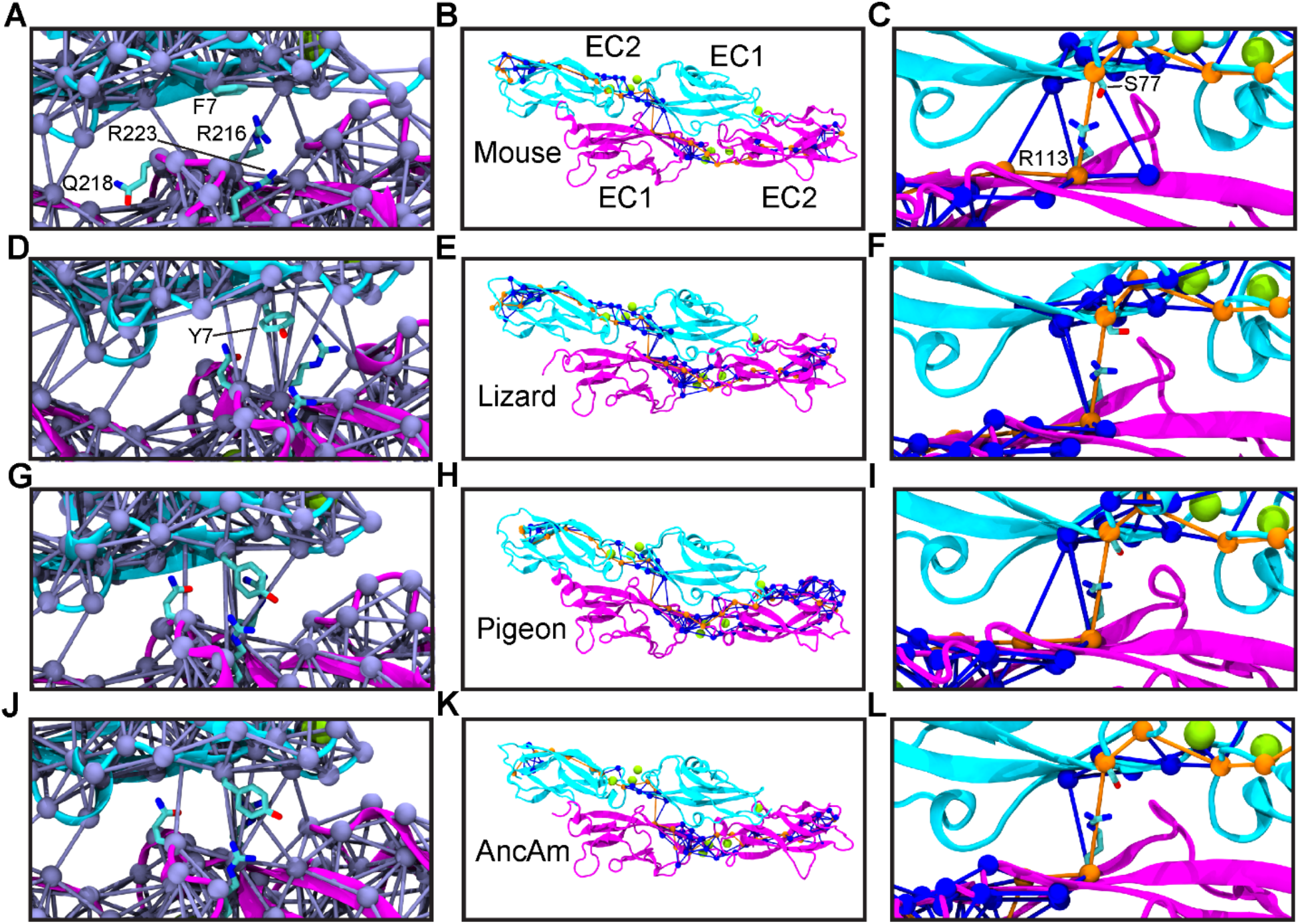
Network analysis of all-atom MD simulations reveals a highly conserved central path of communication between CDH23 and PCDH15 with species-specific changes in the overall network. (*A, D, G* and *J*) The entire dynamic network shows that the mouse (*A*) exhibits fewer connections between the N-terminus of CDH23 and PCDH15 compared to the other three species, likely due to the Y7F substitution in mouse. Analysis from all-atom MD simulations Sim1-4a. (*B, E, H*, and *K*) Optimal (orange) and suboptimal (blue) paths for the species indicated, with the C-termini of each protein selected as source and sink nodes. (*C, F, I* and *L*) While the connectivity of certain suboptimal paths differed between species, most residues along the optimal path were completely conserved, including the point of transfer between CDH23 and PCDH15 that went through R113 and S77 in all four species, consistent with previous results (Hazra et al. 2019).

**FIGURE S5.**
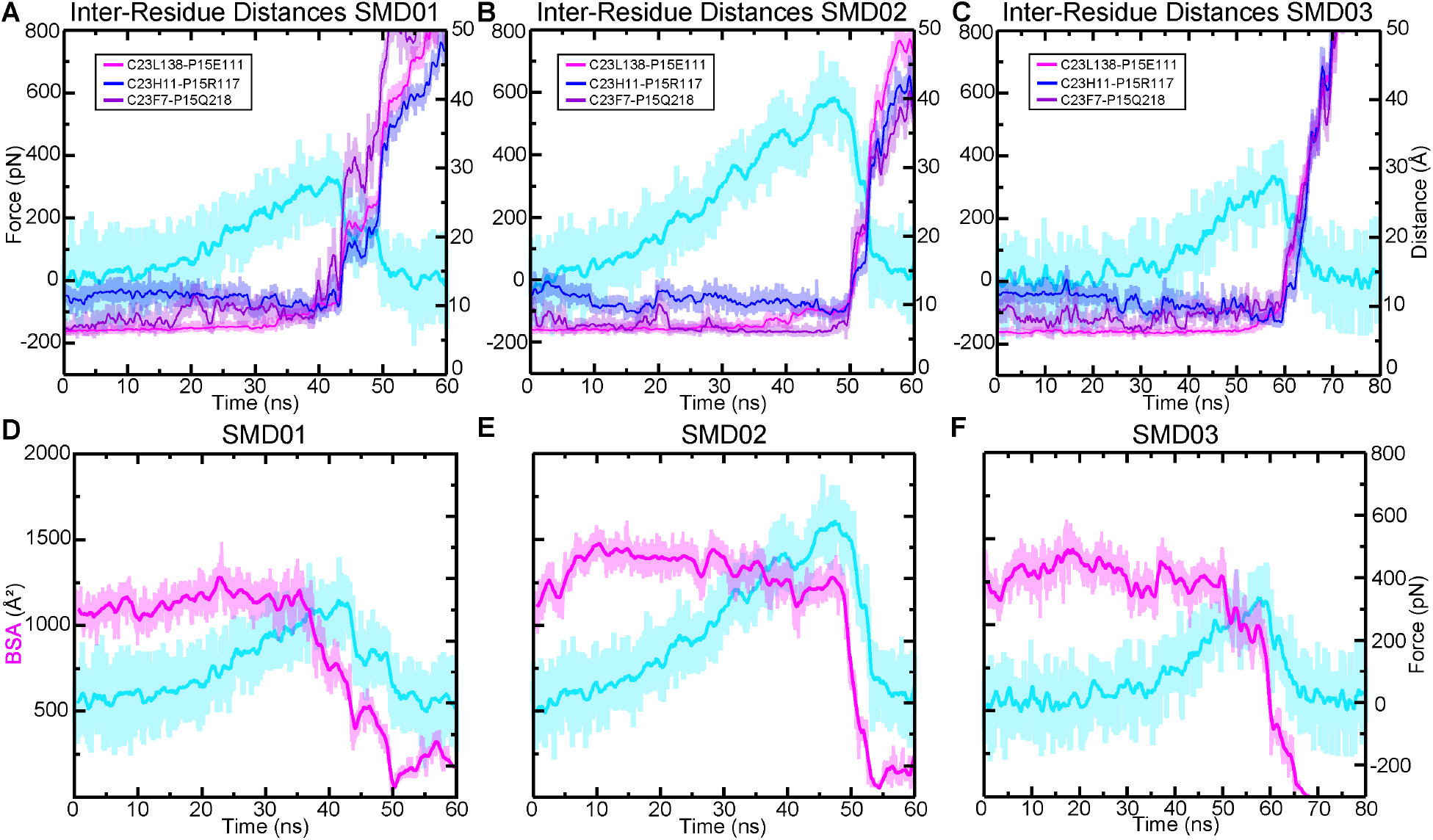
The mouse complex lacks lineage specific interactions and has a weaker interface. (*A-C*) The three interactions (indicated in boxes) in the interface of the mouse complex that differ from the reptilian complexes are shown overlaid with the force profile for the first (*A*), second (*B*), and third (*C*) all-atom SMD simulation (Sim1b-d). The interactions were stable particularly several nanoseconds before and at the force peak in the simulation with a higher force peak (*B*), while in (*C*) and (*A*), the interactions were less stable, particularly before the force peak. (*D-F*) Results from the same three simulations as in (*A-C*), showing the BSA throughout the trajectory. In the simulation with a higher force peak (*E*), the BSA remained higher than in the other two, (*D)* and (*F*), at the force peak.

**FIGURE S6.**
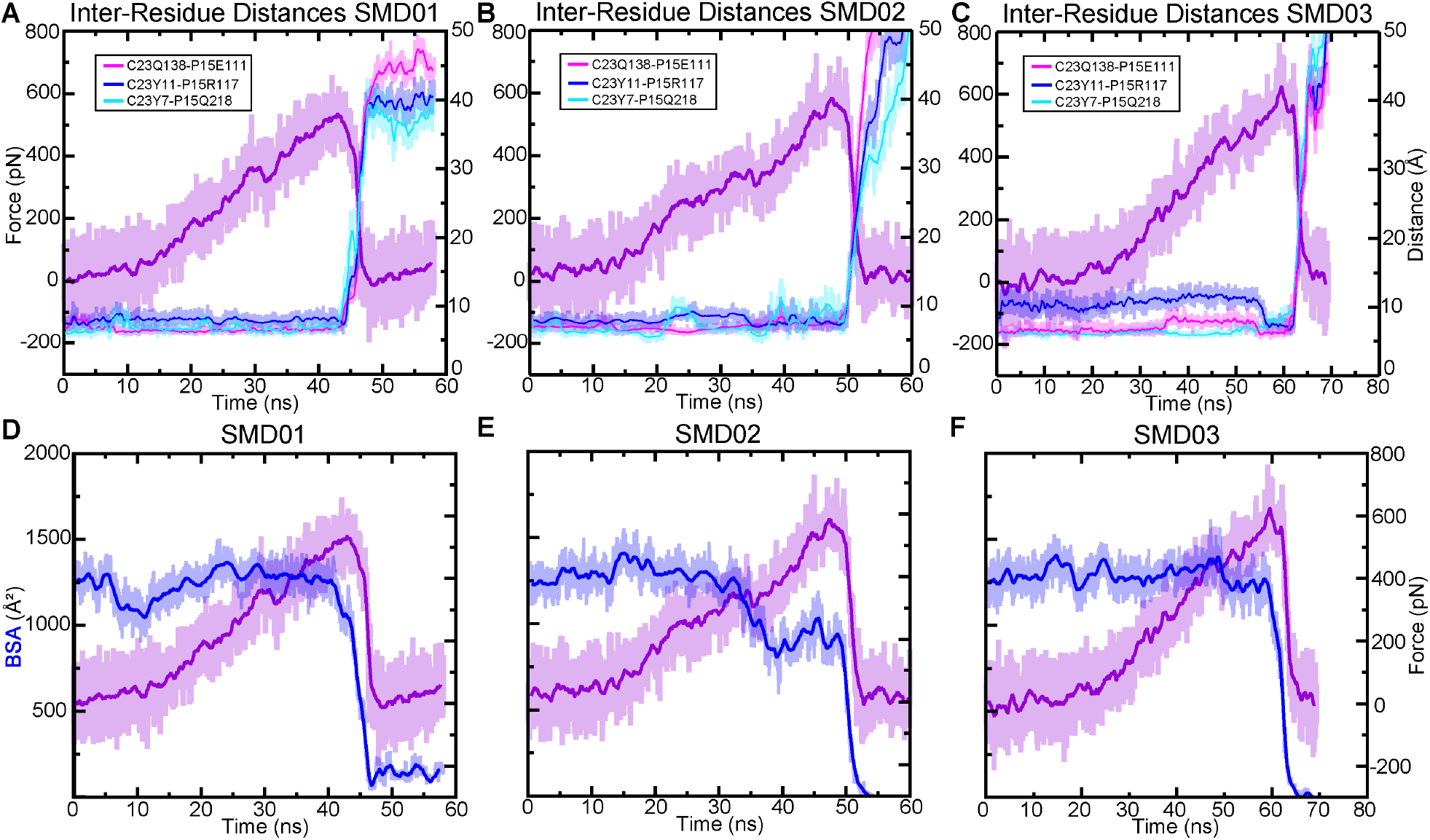
Lineage specific interactions likely contribute to a stronger interface in the lizard. (*A-C*) The three interactions in the lizard interface (indicated in boxes) that differ from the mouse are shown overlaid with the force profile for the first (*A*), second (*B*), and third (*C*) all-atom SMD simulations (Sim2b-d). The interactions were stable particularly several nanoseconds before and at the force peak. This effect is most prominent in the simulation with the highest force peak (*C*). (*D-F*) Results from the same three simulations as in (*A-C*), showing the BSA throughout the trajectory. In the simulation with the highest force peak (*F*), the BSA remained close to its initial value until the force peak.

**FIGURE S7.**
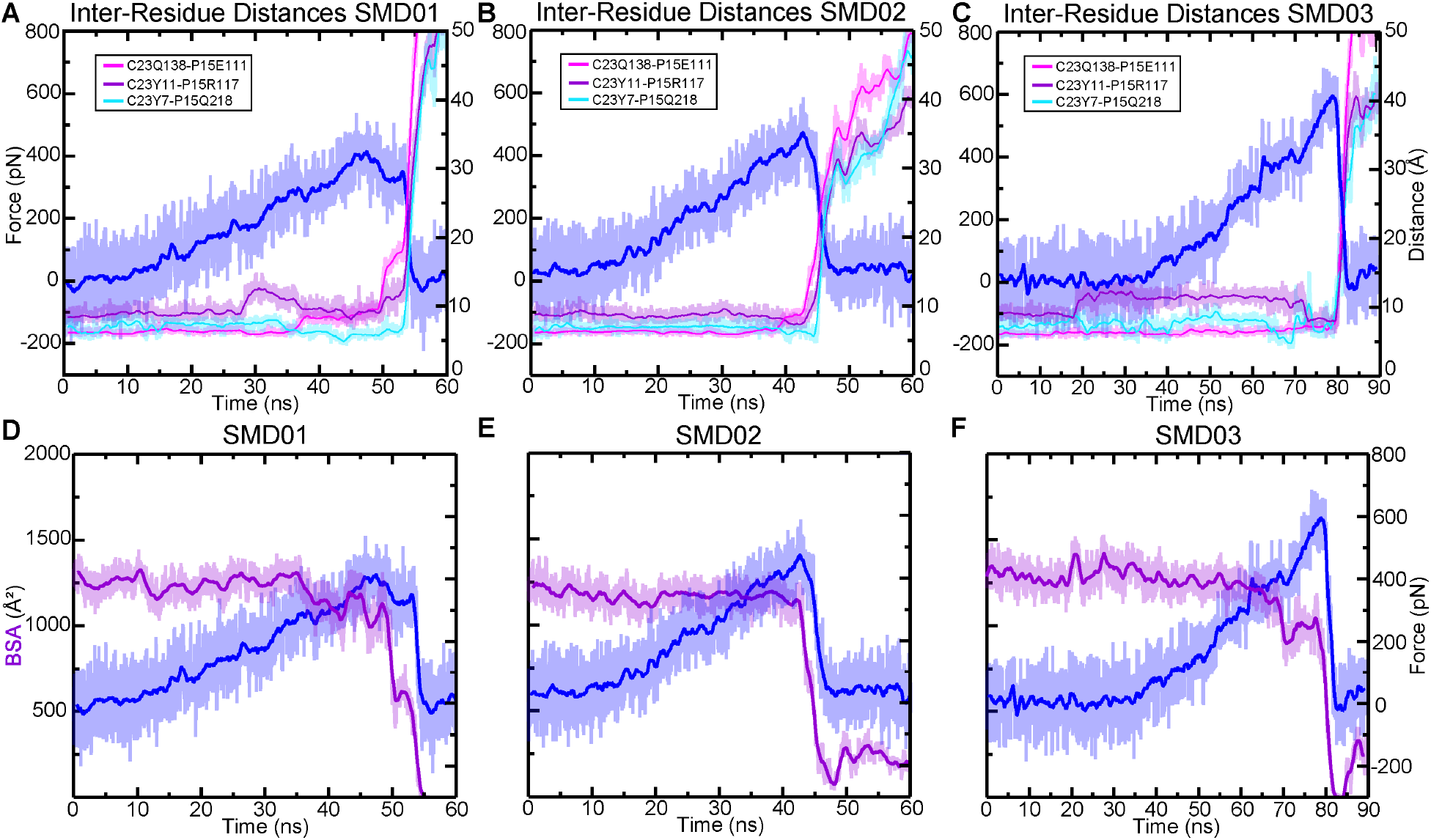
Lineage specific interactions likely contribute to a stronger interface in the AncAm. (*A-C*) The three interactions in the interface (indicated in boxes) in the AncAm that differ from the mouse complexes are shown overlaid with the force profile for the first (*A*), second (*B*), and third (*C*) all-atom SMD simulation (Sim3b-d). The interactions were stable particularly several nanoseconds before and at the force peak in the simulation with higher force peak (*C*), while in (*A*), the interactions are less stable, particularly before the force peak. (*D-F*) Results from the same three simulations as in (*A-C*), showing the BSA throughout the trajectory. In the simulation with a higher force peak (*F*), the BSA dropped before the force peak, but exhibited a sharp increase again at the force peak at ∼70 ns, corresponding to the stabilization of the interactions shown in (*C*).

**FIGURE S8.**
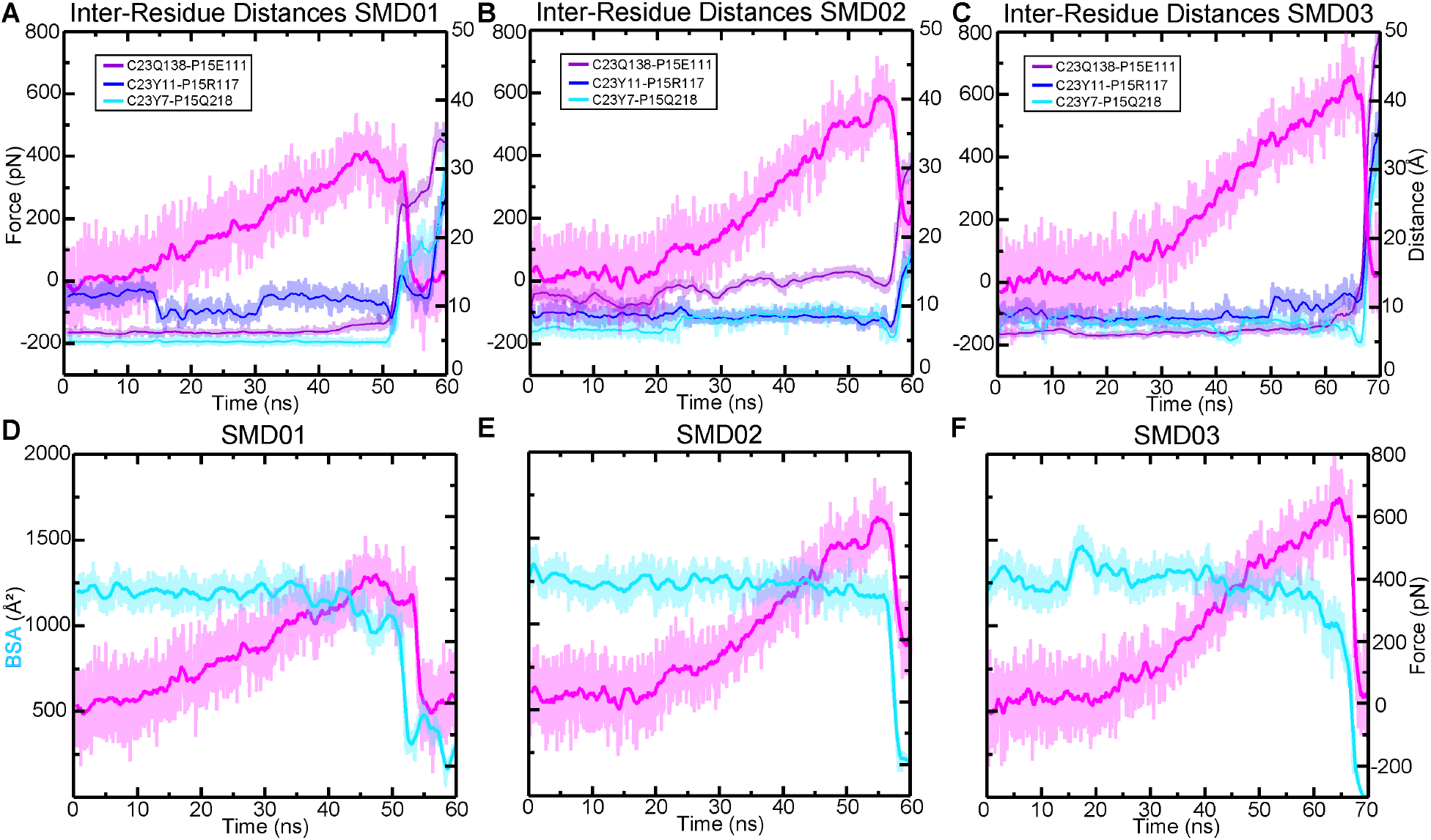
Lineage specific interactions likely contribute to a stronger interface in the pigeon. (*A-C*) The three interactions in the interface in the pigeon (indicated in boxes) that differ from the mouse complexes are shown overlaid with the force profile for the first (*A*), second (*B*), and third (*C*) all-atom SMD simulation (Sim4b-d). The Y11-R117 and Y7-Q218 interactions were stable particularly several nanoseconds before and at the force peak in the simulation with higher force peaks (*B)* and *(C*), while in (*A*), the interactions are less stable, particularly before the force peak, resulting in a lower force. (*D-F*) Results from the same three simulations as in (*A-C*), showing the BSA throughout the trajectory. In the simulation with a higher force peak (*E*), the BSA remained stable until after the force peak.

**FIGURE S9.**
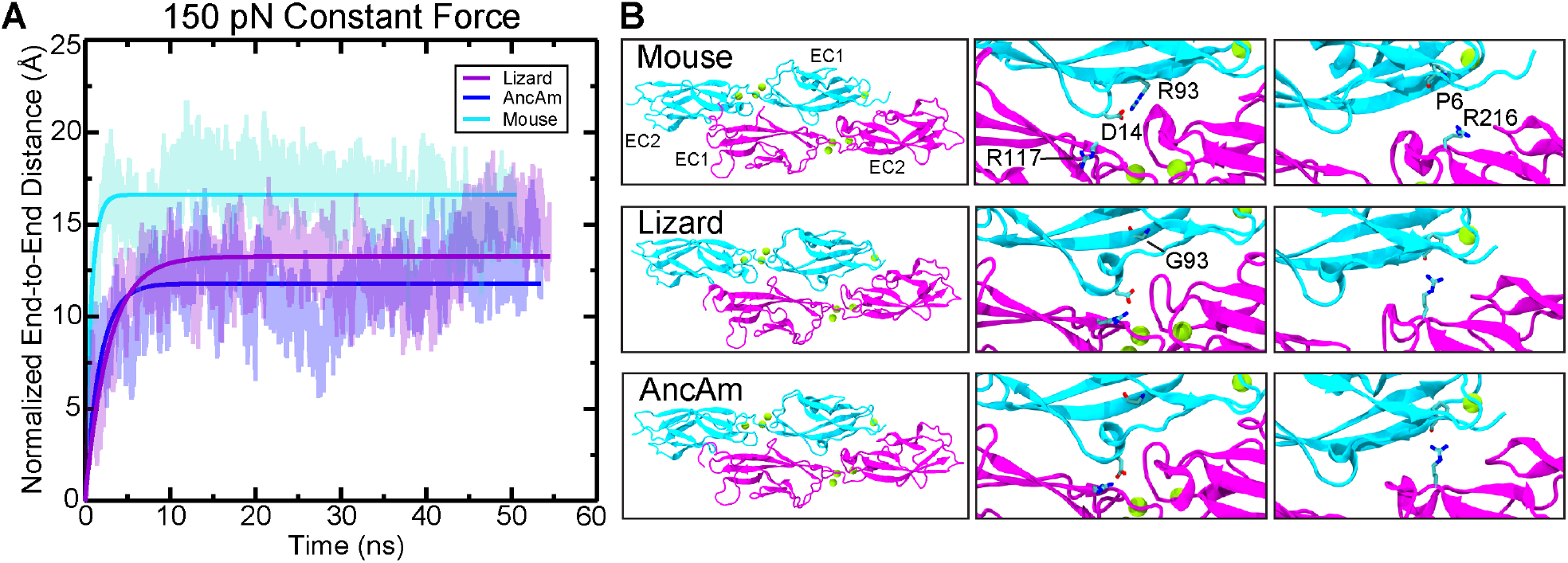
All-atom constant force SMD reveals evolutionary changes in tip-link mechanical properties. (*A*) The mouse exhibits a lower spring constant and damping coefficient (Sim21, *k* = 90.3 pN nm^-1^, γ = 62.4 pN ns nm^-1^) compared to the lizard and AncAm complex (Sim21, *k* = 313.8 pN nm^-1^, γ = 113.1 pN ns nm^-1^ and Sim22, *k* = 127.1 pN nm^-1^, γ = 226.2 pN ns nm^-1^ respectively). (*B*) After application of 150 pN (left panels), the lizard and AncAm exhibit interactions (CDH23 D14-PCDH15 R117 and CDH23 P6-PCDH15 R216) that are not seen in the mouse despite the conservation of the residues directly involved in the interaction (middle and right panels).

**FIGURE S10.**
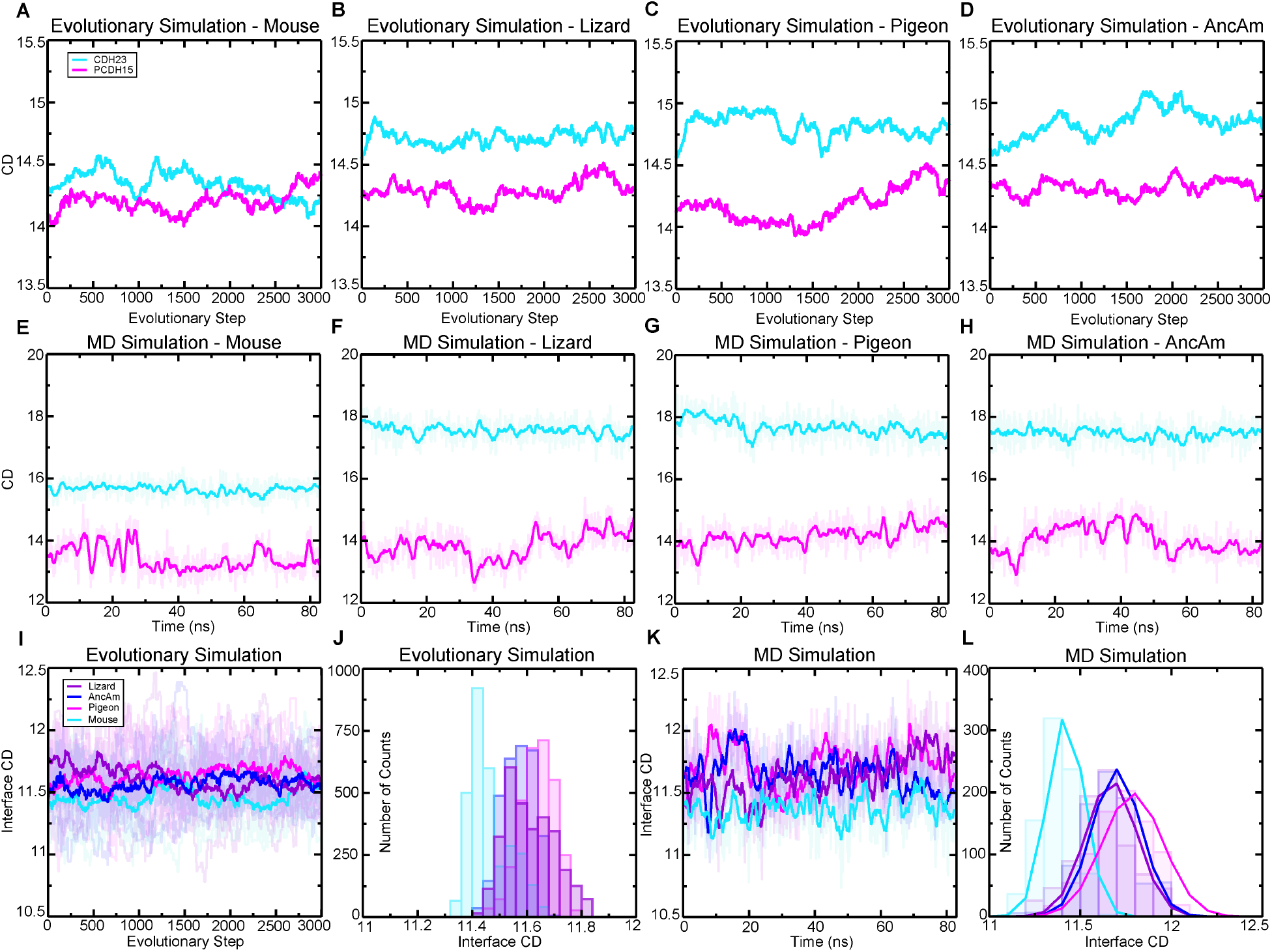
Biophysical properties of CDH23 and PCDH15 show variations that are maintained over nanosecond and evolutionary timescales. (*A-D*) CD of CDH23 and PCDH15 EC1-2 is shown throughout the long-timescale evolutionary simulations. For all species, CDH23 exhibited higher CD throughout the simulations compared to PCDH15, while the mouse exhibited lower CD in CDH23 compared to the other species. (*E-H*) CD of CDH23 and PCDH15 EC1-2 is shown throughout the all-atom MD simulations (Sim1-4a). For all species, CDH23 exhibited higher CD throughout the simulations compared to PCDH15, while the mouse exhibited lower CD in CDH23 compared to the other species. (*I*) Interface CD is shown throughout the long-timescale evolutionary simulations with the average for each species shown in bold. (*J*) The count distributions of the average values from (*I*) are shown for each species, revealing the reduced interface CD for the mouse over evolutionary timescales. (*K*) Interface CD is shown throughout the all-atom MD simulations (Sim1-4a) with the running average for each species shown in bold. (*L*) The count distributions of the average values from (*K*) are shown for each species with a fit to the normal distribution, revealing the reduced interface CD for the mouse during nanosecond time scales.

**FIGURE S11.**
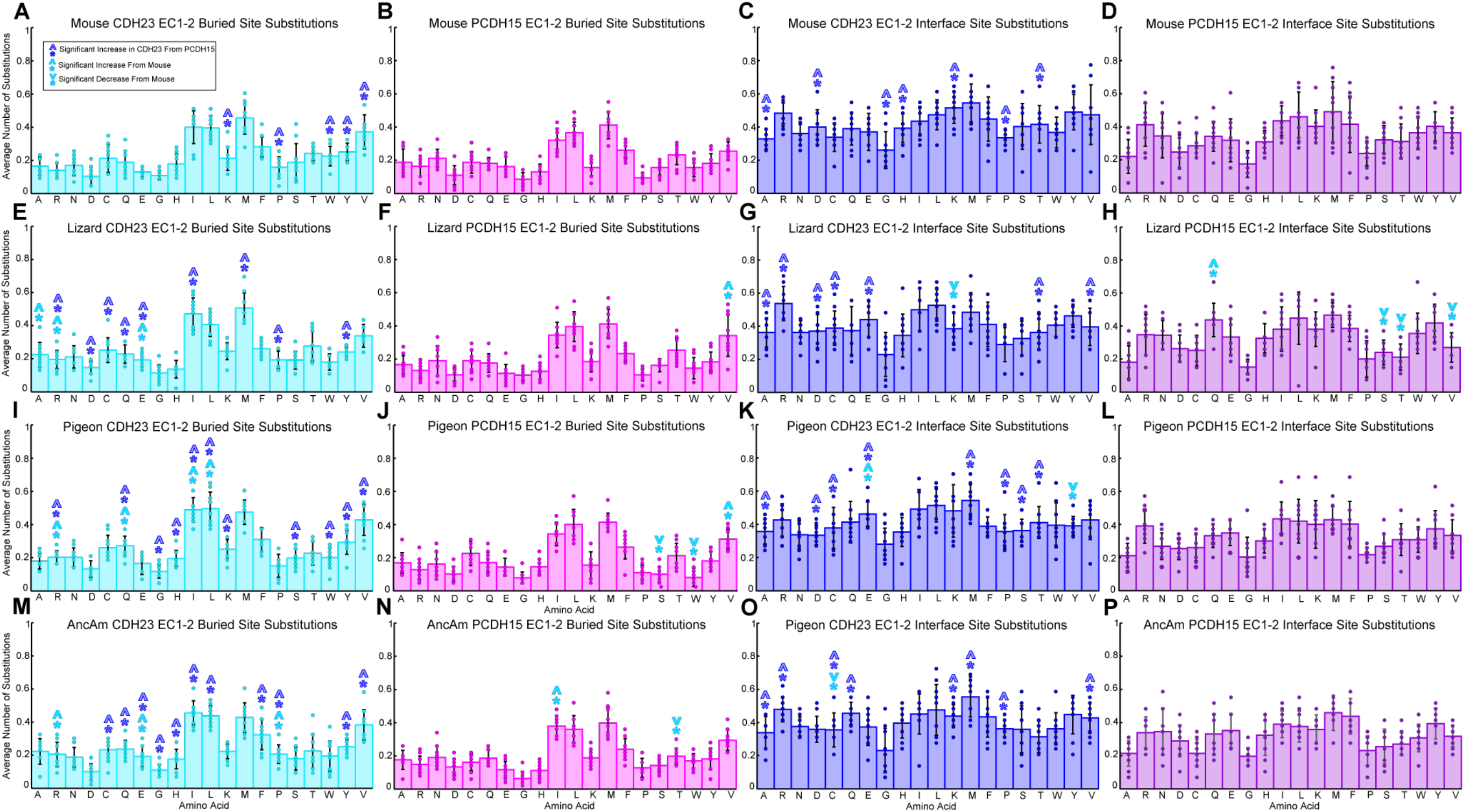
The distribution of amino acids inserted during evolutionary simulations reveals site-wise, protein, and species-specific differences. Average number of insertions per residue, delineated by amino acid, is shown for mouse (*A-D*), lizard (*E-H*), pigeon (*I-L*), and AncAm (*M-P*). Insertions are further broken up between CDH23 and PCDH15 EC1-2, and by the location of the site as indicated. For all species, CDH23 exhibited several significant increases in number of insertions compared to PCDH15 (*A, C, E, G, I, K, M*, and *O*, blue asterisks), while the three non-mammalian species all exhibited significantly increased insertions in CDH23 buried sites compared to the mouse (*A, E, I*, and *M*, cyan asterisks).

---

**TABLE S1.** Evolutionary Simulation Non-Interface Residue Data

| **System** | **Average Number of Fixed Non-Interface Mutations per Residue Long Timescale** | **Average Number of Fixed Non-Interface Mutations per Residue Vertebrate Timescale** | **Average CD From All-Atom MD** | **Average CD From Evolutionary Simulation Long Timescale** | **Average CD From Evolutionary Simulation Vertebrate Timescale** |
| --- | --- | --- | --- | --- | --- |
| *Ac* CDH23 | 7.41 | 0.167 | 17.74 | 14.75 | 14.66 |
| *Ac* PCDH15 | 6.27 | 0.174 | 14.04 | 14.32 | 14.25 |
| *Mm* CDH23 | 7.25 | 0.173 | 15.77 | 14.39 | 14.28 |
| *Mm* PCDH15 | 6.19 | 0.156 | 13.46 | 14.23 | 14.09 |
| *Cl* CDH23 | 7.51 | 0.321 | 17.79 | 14.86 | 14.62 |
| *Cl* PCDH15 | 6.14 | 0.303 | 14.33 | 14.18 | 14.14 |
| AncAm CDH23 | 7.30 | 0.327 | 17.70 | 14.88 | 14.63 |
| AncAm PCDH15 | 6.26 | 0.294 | 14.04 | 14.36 | 14.34 |
| **Total CDH23** | $7.37 \pm 0.02$ | $0.247 \pm 0.089$ | $17.25 \pm 0.99$ | $14.72 \pm 0.23$ | $14.55 \pm 0.18$ |
| **Total PCDH15** | $6.22 \pm 0.01$ | $0.232 \pm 0.078$ | $13.97 \pm 0.36$ | $14.27 \pm 0.08$ | $14.21 \pm 0.11$ |

**TABLE S2.** Evolutionary Simulation Interface Residue Data

| **System** | **Average Number of Fixed Interface Mutations per Residue Long Timescale** | **Average Number of Fixed Interface Mutations per Residue Vertebrate Timescale** | **Average Interface CD From All-Atom MD** | **Average Interface CD From Evolutionary Simulation Long Timescale** | **Average Interface CD From Evolutionary Simulation Vertebrate Timescale** |
| --- | --- | --- | --- | --- | --- |
| *Ac* | 6.97 | 0.155 | 11.67 | 11.62 | 11.71 |
| *Mm* | 7.32 | 0.184 | 11.42 | 11.45 | 11.44 |
| *AncAm* | 7.24 | 0.178 | 11.70 | 11.59 | 11.53 |
| *Cl* | 7.25 | 0.178 | 11.79 | 11.66 | 11.57 |

**TABLE S3.** Evolutionary Simulation Tracking Rejected Mutations Non-Interface Residue Data

| **System** | **Average Number of Fixed Non-Interface Mutations per Residue Tracking Rejected** | **Average CD From Evolutionary Simulation Tracking Rejected** |
| --- | --- | --- |
| *Ac* CDH23 | 0.51 | 14.62 |
| *Ac* PCDH15 | 0.41 | 14.25 |
| *Mm* CDH23 | 0.47 | 14.29 |
| *Mm* PCDH15 | 0.43 | 14.11 |
| *Cl* CDH23 | 0.93 | 14.67 |
| *Cl* PCDH15 | 0.80 | 14.28 |
| *AncAm* CDH23 | 0.92 | 14.74 |
| *AncAm* PCDH15 | 0.78 | 14.34 |
| **Total CDH23** | **0.71 ± 0.25** | **14.58 ± 0.20** |
| **Total PCDH15** | **0.61 ± 0.21** | **14.25 ± 0.10** |

**TABLE S4.**
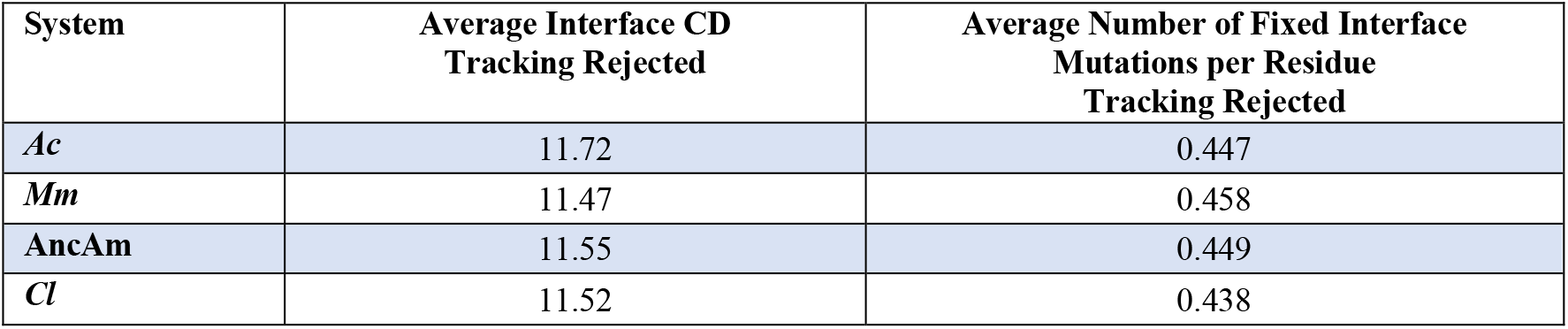
Evolutionary Simulation Tracking Rejected Mutations Interface Residue Data

| System | Average Interface CD<br>Tracking Rejected | Average Number of Fixed Interface<br>Mutations per Residue<br>Tracking Rejected |
| --- | --- | --- |
| *Ac* | 11.72 | 0.447 |
| *Mm* | 11.47 | 0.458 |
| **AncAm** | 11.55 | 0.449 |
| *Cl* | 11.52 | 0.438 |

**TABLE S5.** X-ray Crystallography Experiments

| Construct | Crystallization Conditions | Cryo |
| --- | --- | --- |
| *Ac* CDH23 EC1-2 | 0.1 M MES pH 6.5, 1.1 M CaCl<sub>2</sub> | 25% Glycerol |
| *Cm* CDH23 EC1-2 | 0.1 M TRIS pH 7.7, 1.9 M CaCl<sub>2</sub> |  |
| *AncAm* CDH23 EC1-2 | 0.1 M TRIS pH 7.5, 3 M Sodium Formate |  |
| *AncMam* CDH23 EC1-2 | 0.1 M MES pH 6.5, 1.6 M NaCl |  |

**TABLE S6.**
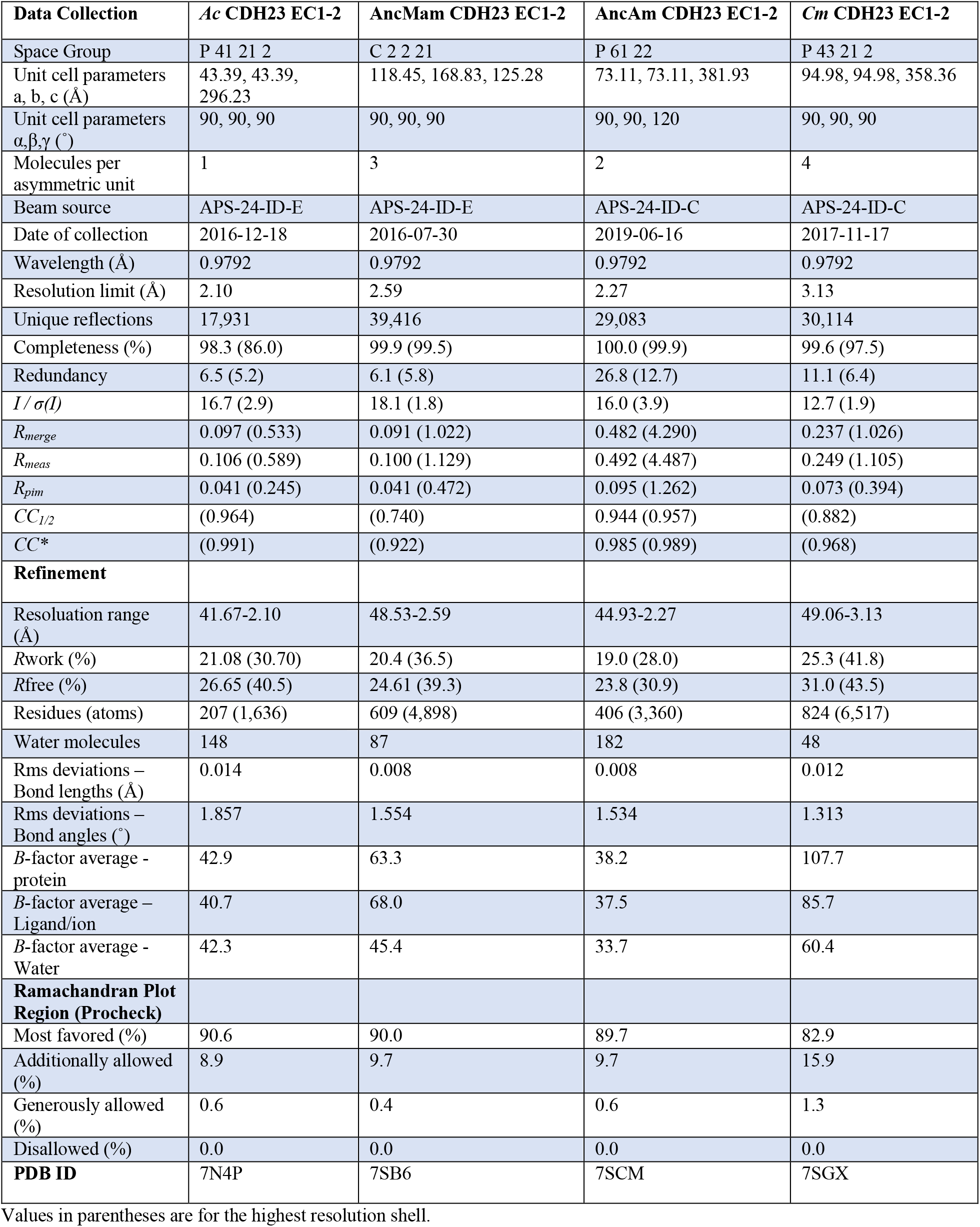
X-ray Crystallography Data Collection and Refinement

| **Data Collection** | ***Ac* CDH23 EC1-2** | ***AncMam* CDH23 EC1-2** | ***AncAm* CDH23 EC1-2** | ***Cm* CDH23 EC1-2** |
| --- | --- | --- | --- | --- |
| Space Group | P 41 21 2 | C 2 2 21 | P 61 22 | P 43 21 2 |
| Unit cell parameters<br>a, b, c (Å) | 43.39, 43.39,<br>296.23 | 118.45, 168.83, 125.28 | 73.11, 73.11, 381.93 | 94.98, 94.98, 358.36 |
| Unit cell parameters<br>$\alpha, \beta, \gamma$ (°) | 90, 90, 90 | 90, 90, 90 | 90, 90, 120 | 90, 90, 90 |
| Molecules per<br>asymmetric unit | 1 | 3 | 2 | 4 |
| Beam source | APS-24-ID-E | APS-24-ID-E | APS-24-ID-C | APS-24-ID-C |
| Date of collection | 2016-12-18 | 2016-07-30 | 2019-06-16 | 2017-11-17 |
| Wavelength (Å) | 0.9792 | 0.9792 | 0.9792 | 0.9792 |
| Resolution limit (Å) | 2.10 | 2.59 | 2.27 | 3.13 |
| Unique reflections | 17,931 | 39,416 | 29,083 | 30,114 |
| Completeness (%) | 98.3 (86.0) | 99.9 (99.5) | 100.0 (99.9) | 99.6 (97.5) |
| Redundancy | 6.5 (5.2) | 6.1 (5.8) | 26.8 (12.7) | 11.1 (6.4) |
| $I / \sigma(I)$ | 16.7 (2.9) | 18.1 (1.8) | 16.0 (3.9) | 12.7 (1.9) |
| $R_{merge}$ | 0.097 (0.533) | 0.091 (1.022) | 0.482 (4.290) | 0.237 (1.026) |
| $R_{meas}$ | 0.106 (0.589) | 0.100 (1.129) | 0.492 (4.487) | 0.249 (1.105) |
| $R_{pim}$ | 0.041 (0.245) | 0.041 (0.472) | 0.095 (1.262) | 0.073 (0.394) |
| $CC_{1/2}$ | (0.964) | (0.740) | 0.944 (0.957) | (0.882) |
| $CC^*$ | (0.991) | (0.922) | 0.985 (0.989) | (0.968) |
| **Refinement** |  |  |  |  |
| Resolution range<br>(Å) | 41.67-2.10 | 48.53-2.59 | 44.93-2.27 | 49.06-3.13 |
| $R_{work}$ (%) | 21.08 (30.70) | 20.4 (36.5) | 19.0 (28.0) | 25.3 (41.8) |
| $R_{free}$ (%) | 26.65 (40.5) | 24.61 (39.3) | 23.8 (30.9) | 31.0 (43.5) |
| Residues (atoms) | 207 (1,636) | 609 (4,898) | 406 (3,360) | 824 (6,517) |
| Water molecules | 148 | 87 | 182 | 48 |
| Rms deviations –<br>Bond lengths (Å) | 0.014 | 0.008 | 0.008 | 0.012 |
| Rms deviations –<br>Bond angles (°) | 1.857 | 1.554 | 1.534 | 1.313 |
| $B$-factor average -<br>protein | 42.9 | 63.3 | 38.2 | 107.7 |
| $B$-factor average –<br>Ligand/ion | 40.7 | 68.0 | 37.5 | 85.7 |
| $B$-factor average -<br>Water | 42.3 | 45.4 | 33.7 | 60.4 |
| **Ramachandran Plot**<br>**Region (Procheck)** |  |  |  |  |
| Most favored (%) | 90.6 | 90.0 | 89.7 | 82.9 |
| Additionally allowed<br>(%) | 8.9 | 9.7 | 9.7 | 15.9 |
| Generously allowed<br>(%) | 0.6 | 0.4 | 0.6 | 1.3 |
| Disallowed (%) | 0.0 | 0.0 | 0.0 | 0.0 |
| **PDB ID** | 7N4P | 7SB6 | 7SCM | 7SGX |

2 Values in parentheses are for the highest resolution shell.

